# Human pan-body age- and sex-specific molecular phenomena inferred from public transcriptome data using machine learning

**DOI:** 10.1101/2023.01.12.523796

**Authors:** Kayla A Johnson, Arjun Krishnan

## Abstract

Age and sex are historically understudied factors in biomedical studies even though many complex traits and diseases vary by these factors in their incidence and presentation. As a result, there are massive gaps in our understanding of genes and molecular mechanisms that underlie sex- and age-associated physiology and disease. Hundreds of thousands of publicly-available human transcriptomes capturing gene expression profiles of tissues across the body and subject to various biomedical and clinical factors present an invaluable, yet untapped, opportunity for bridging these gaps. Here, we present a computational framework that leverages these data to infer genome-wide molecular signatures specific to sex and age groups. As the vast majority of these profiles lack age and sex labels, the core idea of our framework is to use the measured expression data to predict missing age/sex metadata and derive the signatures from the predictive models. We first curated ∼30,000 primary samples associated with age and sex information and profiled using microarray and RNA-seq. Then, we used this dataset to infer sex-biased genes within eleven age groups along the human lifespan and then trained machine learning (ML) models to predict these age groups from gene expression values separately within females and males. Specifically, we trained one-vs-rest logistic regression classifiers with elastic-net regularization to classify transcriptomes into age groups. Dataset-level cross validation shows that these ML classifiers are able to discriminate between age groups in a biologically meaningful way in each sex across technologies. Further, these predictive models capture sex-stratified age-group ‘gene signatures’, i.e., the strength and the direction of importance of genes across the genome for each age group in each sex. Enrichment analysis of these gene signatures with prior gene annotations helped in identifying age- and sex-associated multi-tissue and pan-body molecular phenomena (e.g., general immune response, inflammation, metabolism, hormone response). We developed a web-app (http://mlgenesignatures.org/) to visualize our expression dataset, signatures, and enrichment results to make these easily accessible for interested researchers. Overall, we have presented a path for effectively leveraging massive public omics data collections to investigate the molecular basis of age- and sex-differences in physiology and disease.

**Summary:** Hundreds of thousands of publicly-available human transcriptomes capturing gene expression profiles of tissues across the body and subject to various biomedical and clinical factors present an invaluable, yet untapped, opportunity for studying age and sex. We first curated ∼30,000 primary microarray and RNA-seq samples. Then, we used this dataset to infer sex-biased genes within eleven age groups along the human lifespan and trained machine learning models to predict these age groups from gene expression values separately within females and males. These predictive models capture sex-stratified age-group ‘gene signatures’, i.e., the strength and the direction of importance of every gene in each age group in each sex. Enrichment analysis of these gene signatures with prior gene annotations helped identify age- and sex-associated multi-tissue molecular phenomena. A web-app makes our dataset and results easily visualizable. Overall, we have presented a path for effectively leveraging massive public omics data collections to investigate the molecular basis of age- and sex-differences in physiology and disease.

## Introduction

Most complex traits and diseases have age- and sex-related differences in their incidence and manifestation [1] and yet these factors have been largely under-considered in biomedical and clinical studies in the past [2–5]. Further, often these age- and sex-related differences are intertwined and considering one without the other will produce an incomplete understanding. For example, women have a lower prevalence of stroke before menopause, but afterwards the prevalence exceeds that of men [6]. Similarly, the peak of asthma diagnoses is between the ages of 2 and 8 years old in boys, but incidence is higher for women in adults [7]. As scientific research begins to pay closer attention to age and sex as biological factors, new studies are now beginning to uncover some of the genetic basis that underlies the processes of development and aging with or without considering sex differences in tissue physiology [8], complex traits [9], diseases [10], and treatment responses [11].

These new data alone are not enough to create holistic frameworks capable of helping biologists address questions about female and male tissue biology at specific intervals along the human lifespan (e.g., childhood, adolescence, or old age). A comprehensive understanding of how age and sex influence normal physiology and in turn affect complex traits and diseases is critical to eventually providing individualized medicine for all [12,13]. Related differences can be quite small, producing subtle, easily-overlooked changes. For example, significant sex differentially expressed genes often have small fold changes [12] and genes related to human longevity by GWAS have small effect sizes [14].

An opportunity to investigate these small-yet-widespread influences resides in the hundreds of thousands of publicly-available gene expression profiles generated by hundreds of labs across the world over the past 25 years and stored in databases such as NCBI GEO [15] and EBI ArrayExpress [16,17]. These transcriptomes span multiple tissues, diverse experimental, biomedical, and environmental conditions, and numerous diseases, and have been previously successfully leveraged towards gaining biological insight into molecular mechanisms of complex traits and diseases [18,19]. A few previous studies have used parts of these data to identify sex-associated genes and molecular processes with occasional minor focus on age.

One of the first large-scale endeavors to characterize human sex-biased genes was a 2016 study wherein Mayne and colleagues [20] used differential expression analysis on 22 publicly-available microarray datasets totaling about 2,500 samples from 15 tissues. Previous to that study, one of the largest sex-differential expression studies published was the 2015 study [21] from the GTEx consortium [22] which included sex as one of many biological factors considered in the expression variation between individuals. At the time, only the pilot data had been released, which included 1,641 RNA-seq samples from over 40 tissues in 175 individuals. GTEx consortium data has since grown to over 17,000 samples from 948 individuals and has been used in another handful of sex-differential expression studies. Guo and team [23] used GTEx data in addition to curated datasets from GEO and restricted themselves to healthy samples to determine sex-biased genes in 14 tissues. Gershoni and Pietrokovski [24] published sex-differential expression results using version 6 of GTEX in 2017, and in 2019 the sex-associated gene database (SAGD) resource was published with sex-associated genes (differential expression) from 2,828 samples in 21 different species. Later that year Naqvi et al [25] used GTEx data in conjunction with data from macaque, mouse, rat, and dog to investigate conserved sex-biased expression in 12 tissues. In 2020, Lopes-Ramos and group [26] used the GTEx dataset for sex differential expression analysis and further built regulatory networks for each sample and compared female and male networks in each tissue. Finally, the GTEx consortium released a paper focused entirely on the impact of sex on gene expression in 2020. Each of these studies used publicly-available transcriptomic data, often including GTEx data, and stratified samples by tissue to find differentially expressed genes between sexes. GTEx data skews about ⅔ male, and the Mayne study had a similar ratio of male to female samples. The Guo study, SAGD, and this study are close to 50-50 sex balance. Most studies did not explicitly consider age, but the SAGD considers developmental stage when possible, and the GTEx studies incorporate age as a covariate but is focused on a dataset of mostly adult and older individuals.

Efforts to characterize age-biased genes using human transcriptomic data are also generally stratified by tissue though sometimes commonalities across tissues are investigated. One of the earliest large-scale efforts to study common aging signatures in public expression profiles was an differential expression analysis with 27 microarray datasets from mice, rats, and humans that spanned multiple tissues in 400 samples by Pedro de Magalhães and colleagues [27] in 2007. A few years later Hannum et al [28] used whole blood gene expression data of 488 individuals ranging from 20 to 75 years of age. A large amount of focus in studying age-biased gene expression has continued to center around age prediction [28–33] and the process of aging [27,28,32,34,35] and/or development [36] through differential expression. While sometimes sex is adjusted for in the model, very little has been done to study age in a sex-dependent manner, especially that does not focus on the aging process specifically, though there is evidence that the processes of aging [32,37] and development [38] carry sex-dependent differences.

These important efforts have begun to characterize tissue-specific expression in each sex and in different age groups. However, there are still gaps that need to be addressed. First, most of these studies using public gene expression data are focused on age or sex, sometimes adjusting for the other, but rarely trying to delineate age and sex differences at the same time. Second, the data used in these studies (often from GTEx) skew heavily towards adults and older individuals. Third, investigating both age across all stages of life and sex specificity at the same time on a large scale has yet to be done. We are interested in not just the process of aging, but molecular processes that occur at specific stages along the human lifespan. Integration of a large body of data allows us to pick up on multi-tissue signals without only learning dataset- or tissue-specific signals. The main hurdle to leveraging the hundreds of thousands of publicly-available gene expression profiles towards this purpose is that age and sex metadata is often missing, inconsistent, or disorganized. Especially because age and sex have been historically understudied, the vast majority of these samples are not associated with any age and sex information. Sample descriptions that do contain this information often have it buried in free text and many are annotated with vague labels that are minimally informative and imprecisely defined as, for e.g., ‘old’, ‘adult’, or ‘infant’ (i.e., without the associated age ranges), making it difficult for researchers wishing to reanalyze these datasets.

In this study, we present the largest effort to characterize age- and sex-biased genes across the entire human lifespan from public transcriptomes (**Fig. 1**). First, we manually curated the largest sex- and age-annotated public transcriptome dataset containing nearly 30,000 bulk, primary human microarray and RNA-seq samples from a variety of tissues. Second, to infer pan-body molecular processes impacted by age and sex, we used these transcriptomes and their labels to calculate age-stratified sex-biased gene signatures and sex-stratified age-group-specific gene signatures. Third, using existing gene annotation and association data in various databases, we associated all these age/sex gene signatures to hundreds of biomedical entities including biological processes/pathways, phenotypes, traits, and diseases. We have made the transcriptome dataset, gene signatures, and enrichment results freely available and easily accessible through a web-app (http://mlgenesignatures.org/). Overall, this study presents a framework for leveraging publicly-available omics data collections to study sex-specific health and disease mechanisms in all stages of life.

**Figure 1.**
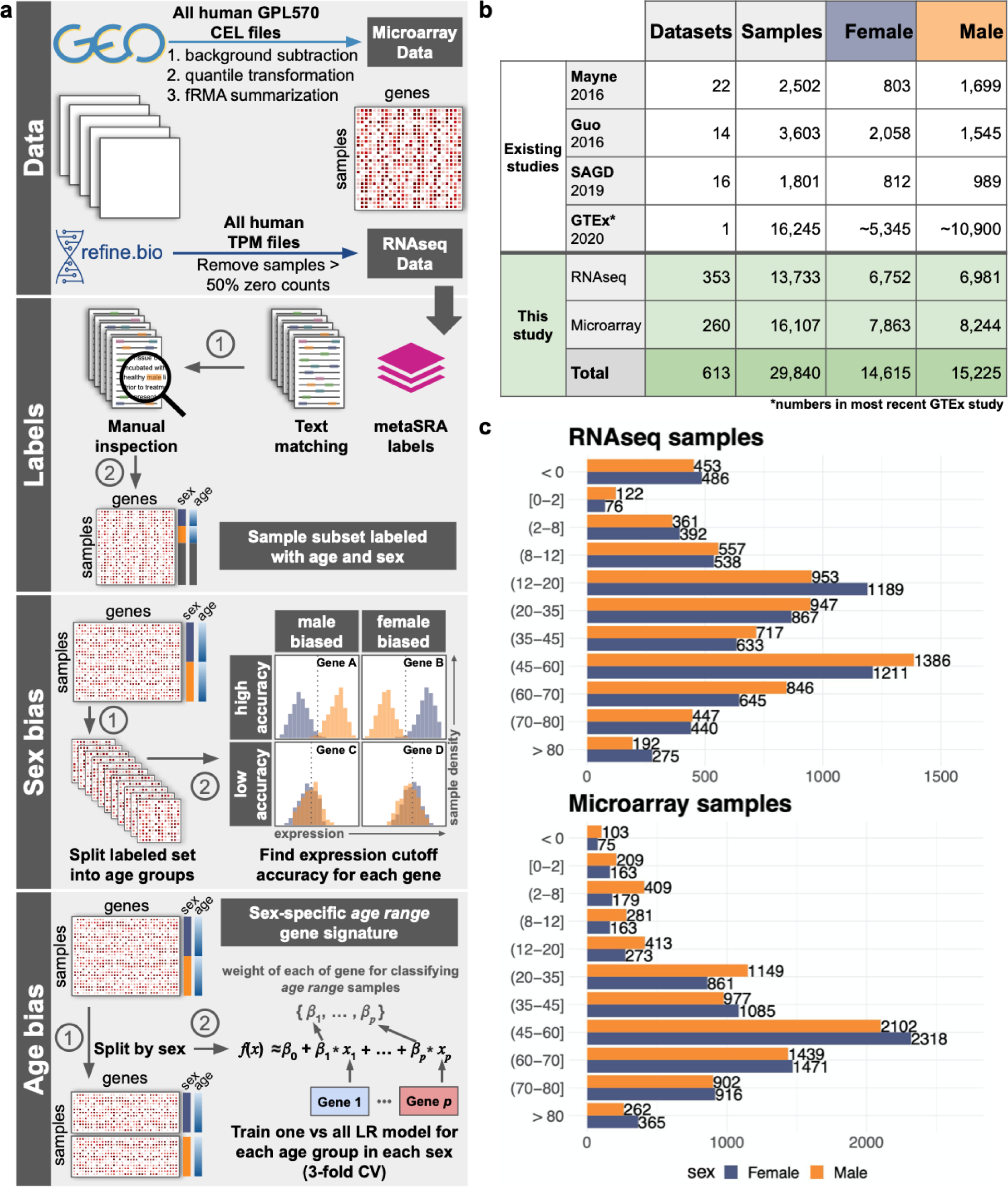
Workflow and data. (**a**) Data was obtained from Gene Expression Omnibus and refine.bio for microarray and RNA-seq, respectively. A combination of text matching and metaSRA labels were used to create a first draft for sex and age labels. These labels were then manually inspected to ensure they were correct and keep only primary human samples. Finally, the labeled data was used to assess sex and age bias in the large, publicly available set. (**b**) The table contains the number of datasets and samples used in some of the largest differential expression studies across sex published in recent years as well as the number of datasets and samples used in this study. The previous studies used only RNA-seq data, while this study uses both microarray and RNA-seq. (**c**) The bar plots show the number of samples used in this study separated by age group (y axis) and sex (bar color).

## Results

### Curating a large dataset of human age- and sex-annotated transcriptomes

To characterize age- and sex-specific gene expression signatures, we first downloaded all available human microarray data in Gene Expression Omnibus (GEO) [39] measured on the same platform and all human RNA-seq data available in refine.bio [40]. We used simple text matching in downloaded sample descriptions from GEO and the Sequence Read Archive (SRA) [41] as well as labels from metaSRA to create a set of transcriptomes associated with age and sex information. We read the sample and experiment descriptions of this entire set of samples to ensure the accuracy of the age and sex labels, as well as to remove any samples that were not bulk, primary human samples.

Our goal with the manual curation step was to assemble a set of samples that would reflect age- and sex-specific molecular mechanisms as faithfully as possible. In addition to improperly assigned age and sex labels, we removed any single cell or single nuclei data, xenografts, microbiome samples, pooled samples, and cell lines. Cell lines were removed due to the tendency of many lines to lose their Y or inactive X chromosome [42], the variability in the ability of cell lines to represent biology of primary cells [43–45], and contamination issues [46]. We divided the lifespan into age groups based on sex hormone levels in each sex across all ages [1]. Through this curation effort, we assembled the largest sex- and age-annotated public bulk transcriptome dataset containing 29,840 samples from human microarray (13,733 samples) and RNA-seq (16,107 samples) technologies (**Fig. 1b**). These samples come from individuals from 11 age groups that span the entire human lifespan, with nearly equal representation from females and males (**Fig. 1c**). Our set has a larger number of samples and datasets in the middle age groups than the youngest and oldest ranges (**Fig. 1c, Fig. S1**) and overall seem to be biased towards samples from blood, brain, small intestine, liver, and lung tissues (**Fig. S2**).

Although tissue bias should be considered, it is worth noting that even when age- or sex-biased genes are determined by restricting samples to a single tissue, cell type composition has a significant effect and will alter results if not controlled for. For example, a handful of studies previously reported breast as the most sex-differentiated tissue [21,24,26], but the most recent GTEx consortium study [47] did not observe this result after controlling for cell type composition. Pellegrino-Coppola and colleagues recently reported a similar importance for cell type correction when determining genes associated with aging from gene expression data [35]. On the task of age prediction, Wang and team found that combining expression data from two tissues reduced the margin of error when predicting the age of GTEx (mostly adult) samples compared to using expression from one tissue [29]. Nonetheless, to investigate the effect of separating out samples by tissue, we repeated some analyses by restricting them to samples from blood only, as it was the most common tissue labeled in our set. However, not enough blood samples were annotated to the fetal (< 0) age group or the oldest age group to include them in the blood-only analyses.

### Age-stratified sex-biased genes

We first used our age- and sex-labeled transcriptome dataset to determine age-stratified sex-biased genes. Independently in microarray and RNA-seq data, we converted the expression of each gene across all samples into z-scores and, for samples within each age group, stepped through the distribution at fixed intervals to find the best expression threshold for that gene to separate ‘Female’ and ‘Male’ samples. Balanced accuracy was used as a metric to determine how well-separated samples were based on each gene’s expression while recording whether the expression was higher in the Female or Male samples (**Fig. 2a**; see ***Methods***). Only 19 genes had a balanced accuracy ≥ 0.8 in at least one age group in either microarray or RNA-seq data and all these genes were on the X or Y chromosome (**Fig. 2c**). As sex differences tend to be quite small and often tissue-specific [12], it is not surprising that all of the strongest sex-biased genes reside in the sex chromosomes. The only Female-biased genes in this set include *XIST*, the major effector of X chromosome inactivation, and *TSIX*, the antisense RNA for *XIST*. The most surprising result is that one gene in this set, *ANOS1,* is both varyingly Male-biased across almost all age groups and is on the X chromosome. Mutations in *ANOS1* have been associated with hypogonadism and Kallmann Syndrome in men [48].

**Figure 2.**
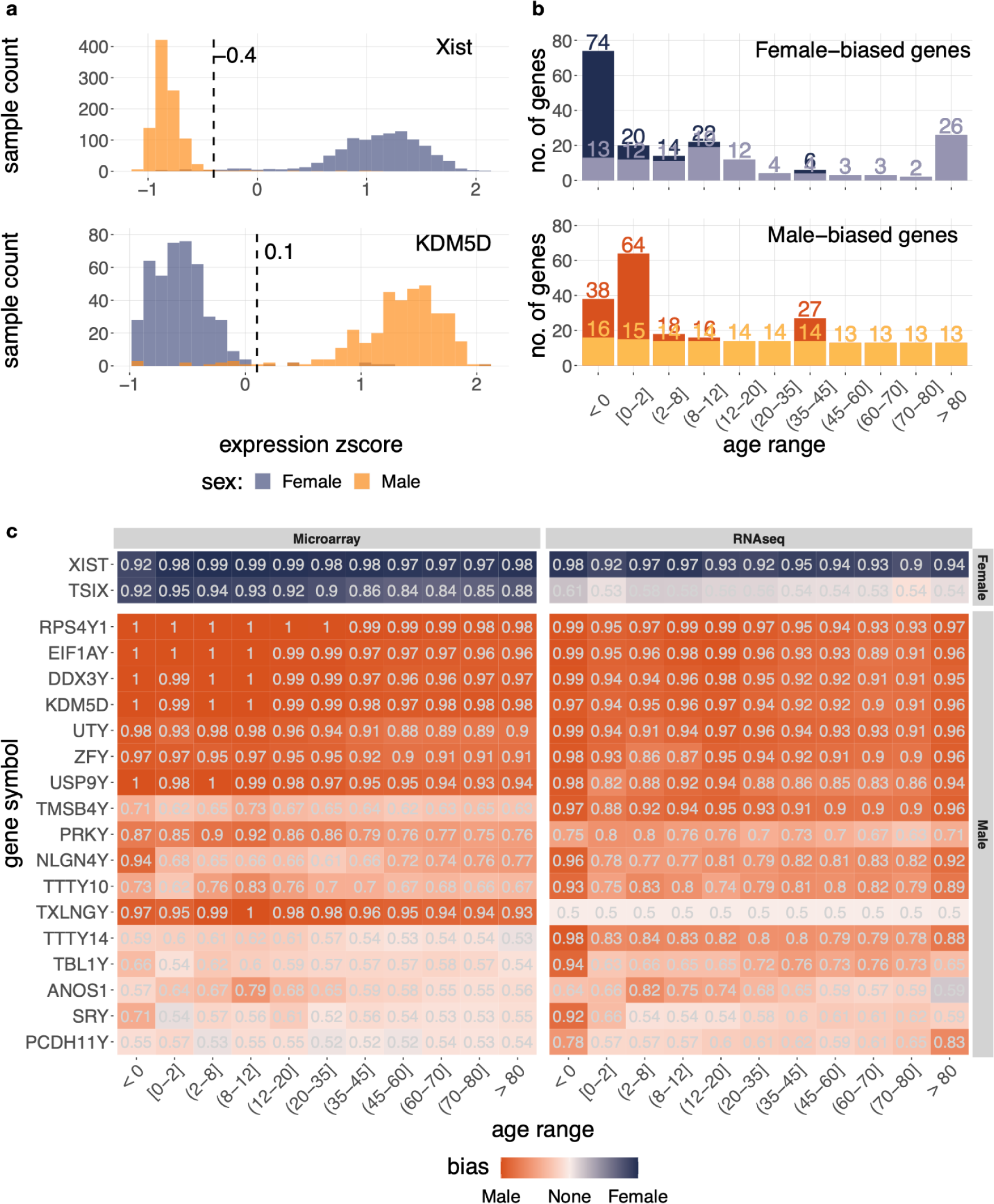
Sex bias of gene expression across age groups. (**a**) Distribution of male and female samples with a given z-scored expression value of Xist (top) and KDM5D (bottom) in different age groups. (**b**) The barplots show the total number of genes with a balanced accuracy of at least 0.65 in one technology and bias agreement between both technologies for Female and Male bias in dark purple and dark orange, respectively. The light purple and light orange bars represent the number of these genes located on the X or Y chromosomes. (**c**) The heatmap displays all genes (x axis) that had a balanced accuracy of at least 0.8 in any age group (y axis) when separating Male and Female microarray or RNA-seq samples.

To ensure that the major signals captured here are not because of tissue-bias across females and males, we repeated this analysis using only samples that came from blood in all but the youngest and oldest age groups due to a low number of samples. Excluding genes reaching the 0.8 threshold only in these age groups from the whole sample set, the blood-only analysis recapitulated all strongly sex-biased genes except *NLGN4Y*, a membrane protein in the neuroligin family (**Fig. S3**). Other sex-biased genes found in blood samples were restricted to a single age group each.

We then examined genes with weaker sex bias, but reasonably consistent across technologies. These were genes biased in the same direction in both RNA-seq and microarray data, but only reached a balanced accuracy of 0.65 in one of the technologies (**Fig. 2b**). No Y chromosome genes that did not reach a balanced accuracy of 0.8 in at least one age group were added in this set, but several more genes from the X chromosome were added, including 4 more genes that were Male-biased (*DUSP9*, *CD99*, *THOC2*, *SMARCA1*) in some of the younger age groups. The only Female-biased X chromosome gene common across all age groups was *KDM6A*, a lysine demethylase. The next most common Female-biased X-chromosome genes across age groups were *EIF1AX*, *PUDP*, and *ZFX*. These were all common to the seven youngest age groups along with the oldest age group. Interestingly, general sex-biased expression seems to taper off as age increases (**Fig. 2b**), but the oldest age group contained a particularly high number of Female-biased genes (26), all on the X chromosome. These genes were not enriched for any particular function. The youngest age groups also show the most sex bias in autosomal genes; the four oldest age groups showed no sex-biased autosomal genes.

### Age group prediction stratified by sex

Next, for each sex, we used our dataset to scan for genes that were able to differentiate between age groups to find sex-stratified age-biased genes. We set this task up as a supervised machine learning problem and trained logistic regression models to distinguish one age group from all others, independently for each sex, in RNA-seq (**Fig. 3**) and microarray (**Fig. S4**) data separately. We used an elastic-net penalty to encourage sparsity while balancing the contributions of correlated genes. Entire datasets were always kept within a fold to avoid rewarding the model for learning study-specific signals (**Fig. S5**). Across all age groups, ML models based on RNA-seq use less genes to predict sample age range (**Fig. 3a, Fig. S4a,c**). In both technologies, the middle age groups are harder to separate from other age groups using gene expression, as evidenced by the higher number of genes required by the model and slightly lower performance compared to the very youngest and oldest age group models (**Fig. 3** and **S4**).

**Figure 3.**
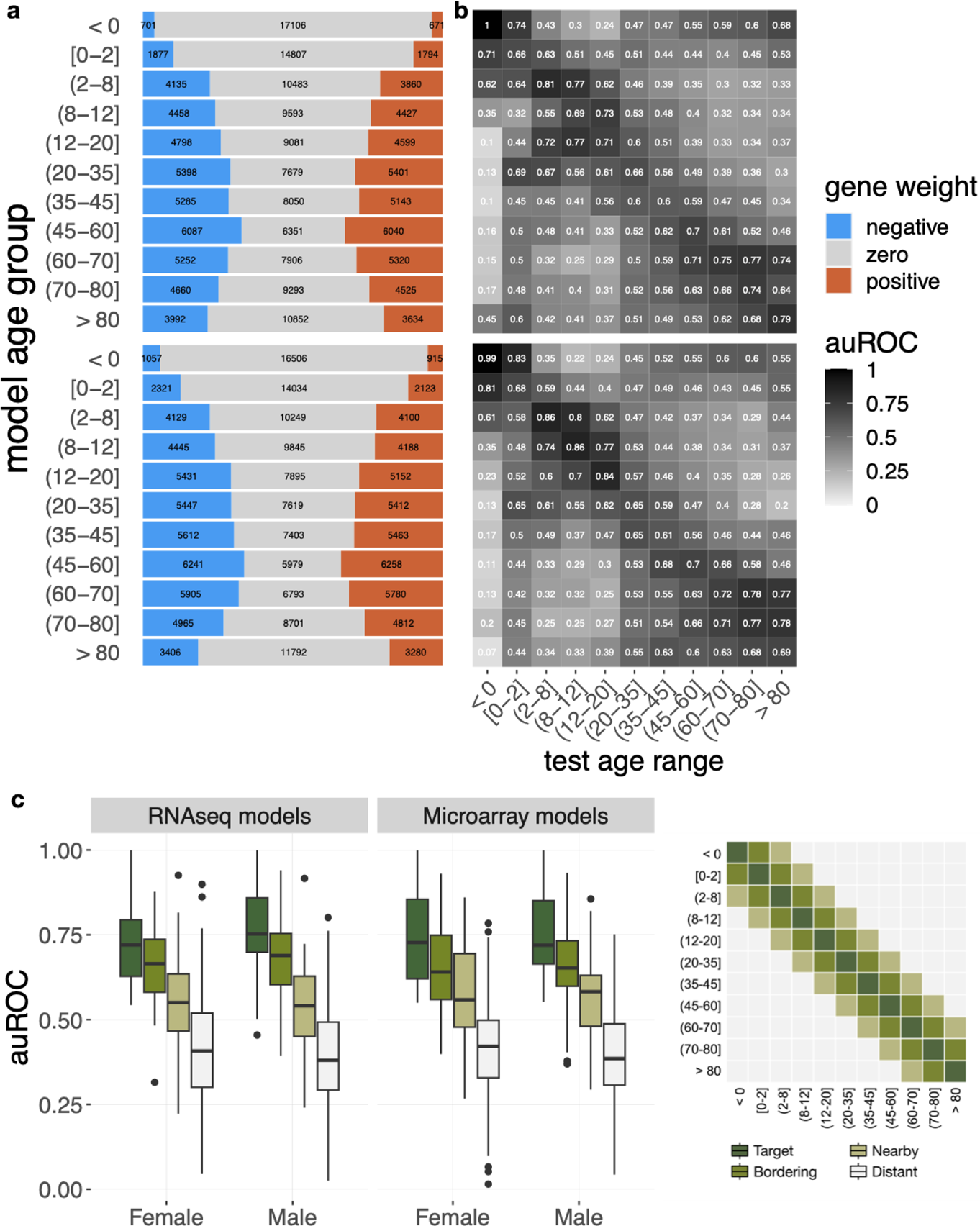
Size and performance of age group prediction models. (**a**) The stacked barplots show the distribution of positive, zero, and negative weights for the model with the median number of positives across the three folds for each age group in RNA-seq for Females (top) and Males (bottom). (**b**) The heatmaps contain average auROC of RNA-seq models trained on the age group labeled in the rows when evaluated using the samples in the age group labeled in the column as positive examples for Females (top) and Males (bottom). (**c**) Boxplot of auROC of all RNA-seq and microarray Female and Male models when considering target, bordering, nearby, and distant age groups as positive examples (key on right).

In addition to measuring the performance of each model on the age group it was trained to classify, we also evaluated it on samples from each of the other age groups (**Fig. 3b, Fig. S4b,d**). The heatmaps in **Figure 3** and **Figure S4** contain the average area under the Receiver Operator Characteristic curve (auROC) across 3 folds for each age group model trained on a specific age group (rows) and evaluated as if the age group in the column were the positive examples. We chose to display auROC as it can be easily interpreted as a probability. Thus, the value in each cell of a heatmap is the probability that a randomly selected sample from the test age group (column) would be ranked higher than a randomly selected sample from any other age group by the age group model in the row. However, auROC is not the best measure of performance when there is a high imbalance of positive and negative examples, as we have in this age-group classification task. So, we also include heatmaps of the performance measured with log_2_(auPRC/prior) in the supplement (**Fig. S6-9**). This metric accounts for class imbalance and emphasizes the accuracy of top-ranked positive samples. Nevertheless, evaluation results with this metric largely agrees with those shown by auROC.

Overall, the classifiers show good performance across age groups in both RNA-seq and microarray samples, with more difficulty in the middle age groups (**Fig. 3b, Fig. S4b,d, Fig. S6-9**). We also see poorer performance in the [0–2] age group RNA-seq models, where we have one of the lowest number of positive examples (**Fig. 3b**, **Fig. 1c**). In general, however, the number of training positives does not correlate with higher performance (**Fig. S10**). For instance, the (45–60] age group in both sexes has the worst prediction performance with microarray samples despite having the highest number of samples (**Fig. S4b,d,** **Fig. 1c**).

We repeated this analysis using only blood samples for training and evaluation using three test sets, but reusing some training data in all folds because the lower number of samples made it impossible to perform a strict three-fold cross validation for some age groups. Even with this adjustment, we still were not able to include the youngest and oldest age groups (**Fig. S11**). Overall, the blood-only models used less genes to classify samples (more genes had zero weight) than the models using all tissues, but had poorer prediction performance in most age groups (**Fig. S12-17**). This result suggests that including data from multiple tissues may improve the age signal-to-noise ratio (see ***Discussion***).

To check consistency between folds and similarity between models in different age groups in both sexes and technologies, we calculated the cosine similarity of the model weights across all genes (**Fig. S18–22**). As expected, invariably, models trained on the same age group, sex, and technology are more similar to each other than to models that differ in any of those factors. For the youngest age groups, especially < 0, we observe the models to be similar even across sex and technologies. This observation combined with the high performance of these models indicates that fetal gene expression is robust and very distinct from all other age groups. Conversely, most sex-stratified age group models are not similar across RNA-seq and microarray, indicating a substantial technology effect.

Finally, the cross-age-group evaluations (**Fig. 3** and **S4**) also demonstrate that the age group models capture the chronological relationships between age groups. We quantified this pattern more precisely by dividing all age groups into four categories with reference to each model. The ‘target’ age group is the one that the model was trained on, ‘bordering’ age groups are those directly before or after the target, and ‘nearby’ age groups are those that are one-removed from the target. Every other age group is marked as ‘distant’. As we train a one-vs-rest model for each age group, the model is designed to separate target (positive) samples from non-target (negative) samples and does not have any external information about the chronological relationships between age groups. Despite this setting, we observe that age group models assign higher probabilities to samples from neighboring age groups and lower probabilities to those from distant age groups (**Fig. 3c**). This trend holds in both sexes for RNA-seq and microarray models and is strong evidence for our models capturing biologically-relevant age signals. A similar trend can be observed in the blood age group model performances, but it is not quite as strong, especially in Females (**Fig. S23**).

### Sex-stratified age-biased genes

We defined sex-stratified age-biased genes per age group by choosing genes that were assigned a positive weight in the corresponding model in at least five folds across the six folds between the microarray and RNA-seq models, with a non-negative weight in the remaining fold, if any. The middle age groups have a higher number of age-biased genes than other age groups (**Fig. 4a,b**, (**Fig. 3a, Fig. S4a,c**)). Across all age groups in both sexes, the number of age-biased genes from each chromosome tends to correlate with the total number of genes on the chromosome (**Fig. 4a,b**). Similar trends are present in negatively-weighted genes from models in each sex (**Fig. S24**).

**Figure 4.**
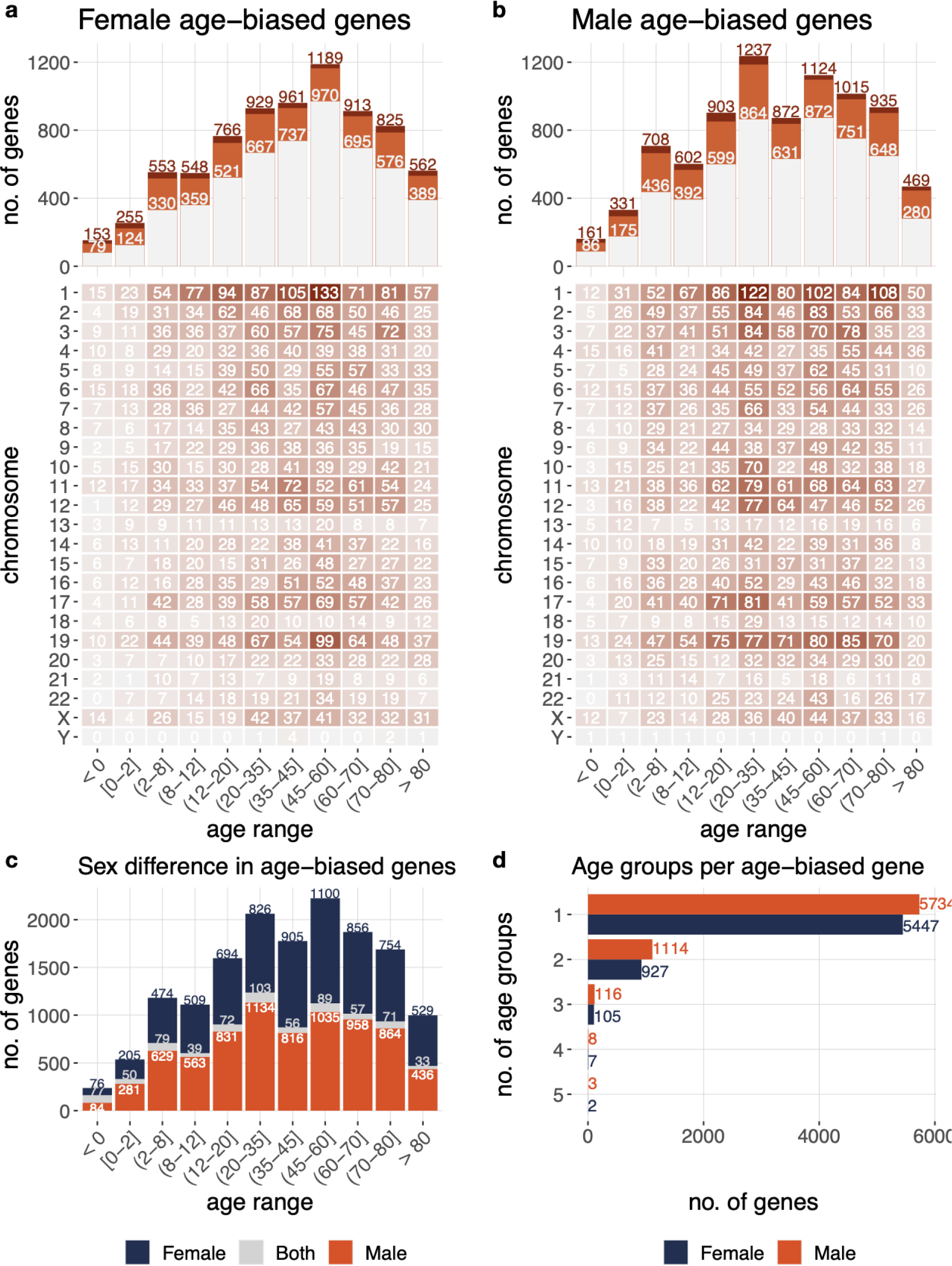
Number of age-biased genes across age groups in each sex. The top two barplots show the total number of age-biased genes per age group in (**a**) Females and (**b**) Males. The tables display the number of age-biased genes on each chromosome per age group in (**a**) Females and (**b**) Males. (**c**) The number of age-biased genes per age group that are age-biased in Females only (purple), Males only (orange), or in both sexes (light gray). (**d**) The number of age groups per sex that each age-biased gene is highly weighted in.

In total, across all age groups, 6,488 genes are age-biased in Female models and 6,975 genes are age-biased in Male models, but only 2,838 of those genes are common between them. Even within age groups, the overlap of age-biased genes between sexes is quite low (**Fig. 4c**) and the vast majority of these age-biased genes are biased in only one age group per sex (**Fig. 4d**). In each sex, about 1,000 genes are age-biased in two age groups, around 100 in three age groups, and about 10 total across four or five age groups (**Fig. 4d**). Taken together with the good performance of our prediction models, these results suggest that each age group has a distinct expression signature and that development and aging processes have sex-specific differences detectable in large datasets.

### Enriched experimental genesets in age-stratified sex signatures

All the analyses presented above have resulted in several genome-wide gene signatures associated with sex and age. To enable us and other researchers efficiently interpret (and search) these signatures in relation to various biological contexts, we associated these signatures to coherent genesets annotated to biological processes [49], traits [50], diseases [51,52], phenotypes [53], and cell types [54]. First, we defined age-stratified sex gene signatures as genes across the genome along with their signed normalized balanced accuracy scores, which indicates the extent to which each gene was able to separate Female from Male samples (or vice versa) in a particular age group (See ***Methods***). We then used a permutation test to estimate the strength and direction of association (*i.e.*, ‘enrichment’) of each geneset with each signature (see ***Methods***).

We applied this approach to compare the sex gene signatures calculated in our study and those from previous studies — Guo et al (2016) [23], SAGD (2019) [55], and GTEx (2020) [47] — estimated using differential expression analysis in multiple tissues. Overall, the agreement between studies is very high. The only prominent disagreement occurs with GTEx signatures, especially in our [0–2] sex signature (**Fig. S25**). However, there are no samples from children in GTEx, which makes it unsuitable for capturing what sex-biased expression should look like in children no older than 2. All the other partial and minor disagreements likely point to sex biases in specific age groups found in our analysis that were not seen in previous analyses that did not stratify data by age. However, the allround agreement suggests that gene signatures we estimate reflect patterns of sex differences in expression across various tissues.

Application of the geneset enrichment approach to genesets from various sources resulted in hundreds of biological contexts associated with sex across age group. Notable among them is the pan-body phenomenon of immune response that plays a central role in autoimmune diseases more prevalent in females [56,57]. We found the large majority of immune response-related (**Fig. 5a**) and viral-related (**Fig. 5b**) biological processes to be Female-biased across most age groups. Notable exceptions include the < 0 and (2-8] age groups. A very similar trend can be observed with the enrichment of immune diseases (**Fig. 5c**) and immune phenotypes (**Fig. 5g**).

**Figure 5.**
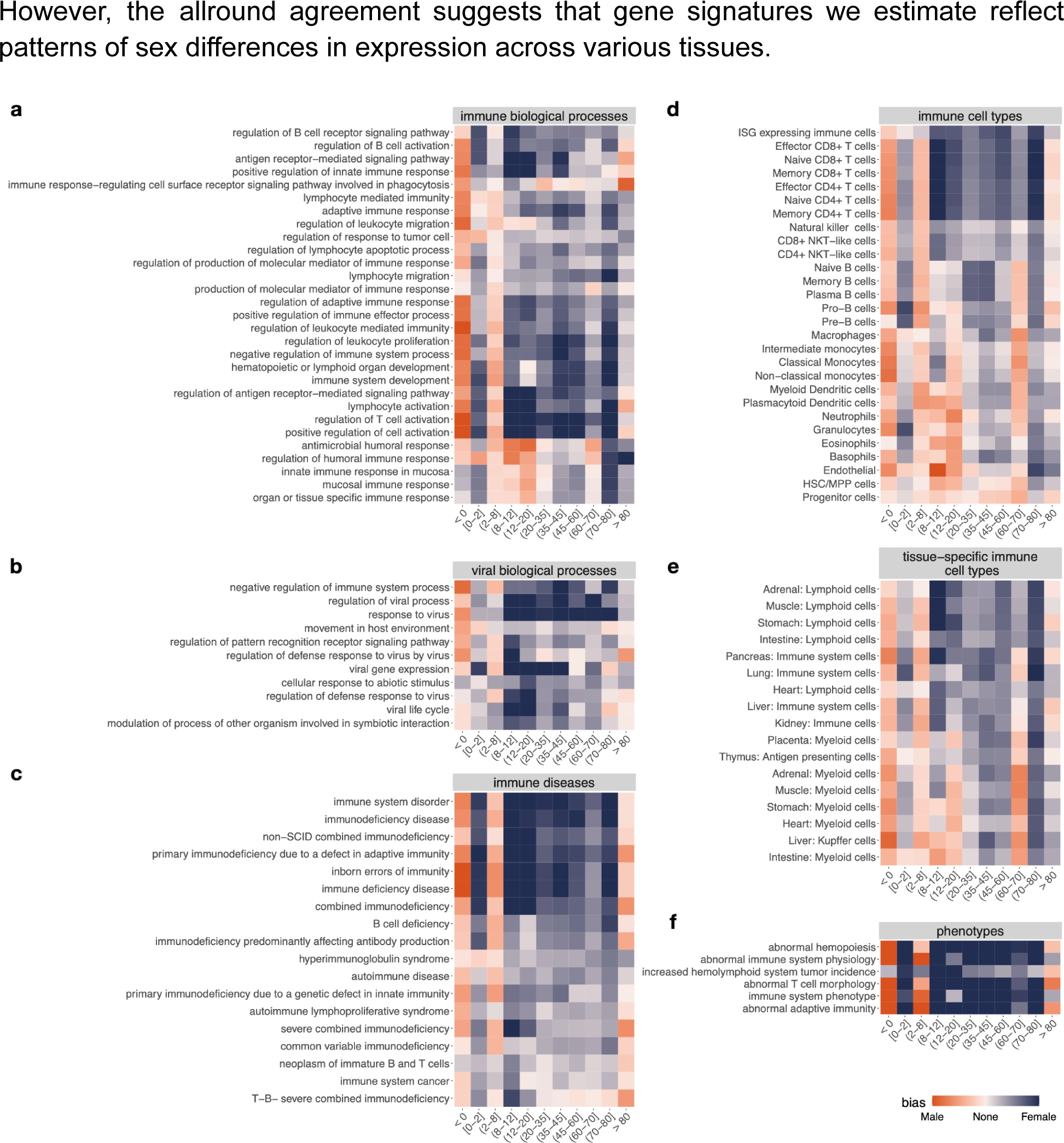
Enrichment of experimentally-derived genes sets from in our age-stratified sex signatures. Female- and Male-bias enrichment of experimentally-derived gene sets. Heatmaps show enrichment scores for (**a**) a representative (See ***Methods***) set of immune-related GO biological processes, (**b**) a representative (See ***Methods***) set of viral-related GO biological processes, (**c**) immune-related diseases, (**d**) immune cell type marker genes, (**e**) tissue-specific immune cell type marker genes, (**f**) Schwann cell marker genes, and (**e**) immune-related phenotypes.

Consistent with previous studies [58,59], the marker genes of B cells, T cells, and lymphoid cells tend to be Female-biased in our sex signatures (**Fig. 5d, e**), again with the exception of the < 0 and (2-8] age groups, and the (60-70] age group for B cells. We also observe that myeloid cells tend to have Male-biased enrichment in the younger age groups (consistent with a previous study [59]) and Female-biased enrichment in older age groups.

Further, we find many metabolic processes to be Male-biased in our age-stratified sex signatures, consistent with findings from a previous study [23] (**Fig. S26**). Together, these enrichment results indicate the potential utility of our signatures to investigate sex differences of multi-tissue processes such as immune response and metabolic processes, along with corresponding disease mechanisms in different stages of the human lifespan.

### Enriched experimental genesets in sex-stratified age group signatures

In addition to being valuable for making age group prediction of new samples, our age group models can be biologically interpreted. To do so, we first defined sex-stratified age-group gene signatures as genes across the genome along with their model coefficients. Here, the genes with positive and negative coefficients in a model correspond to genes whose relative high and low expression, respectively, are characteristic to the pertinent age group (see ***Methods***). Then, we used the same permutation-based geneset enrichment strategy as above to associate hundreds of biological contexts — biological processes, traits, diseases, phenotypes, and cell types — to these signatures (see ***Methods*** for details).

We began our investigation of these results by focusing on biological processes that show enrichment strongly correlated with increasing (**Fig. 6a,b**) or decreasing (**Fig. 6c,d**) age. Apoptosis-related processes are associated with relatively low or non-expression in young age groups but show increased association with older age groups in age signatures in both sexes, especially in Females, reminiscent of observed aging processes [60]. In Males, positive regulation of NF-kappaB signaling and negative regulation of lymphocyte migration also increase in association with age, consistent with other studies [59,61].

**Figure 6.**
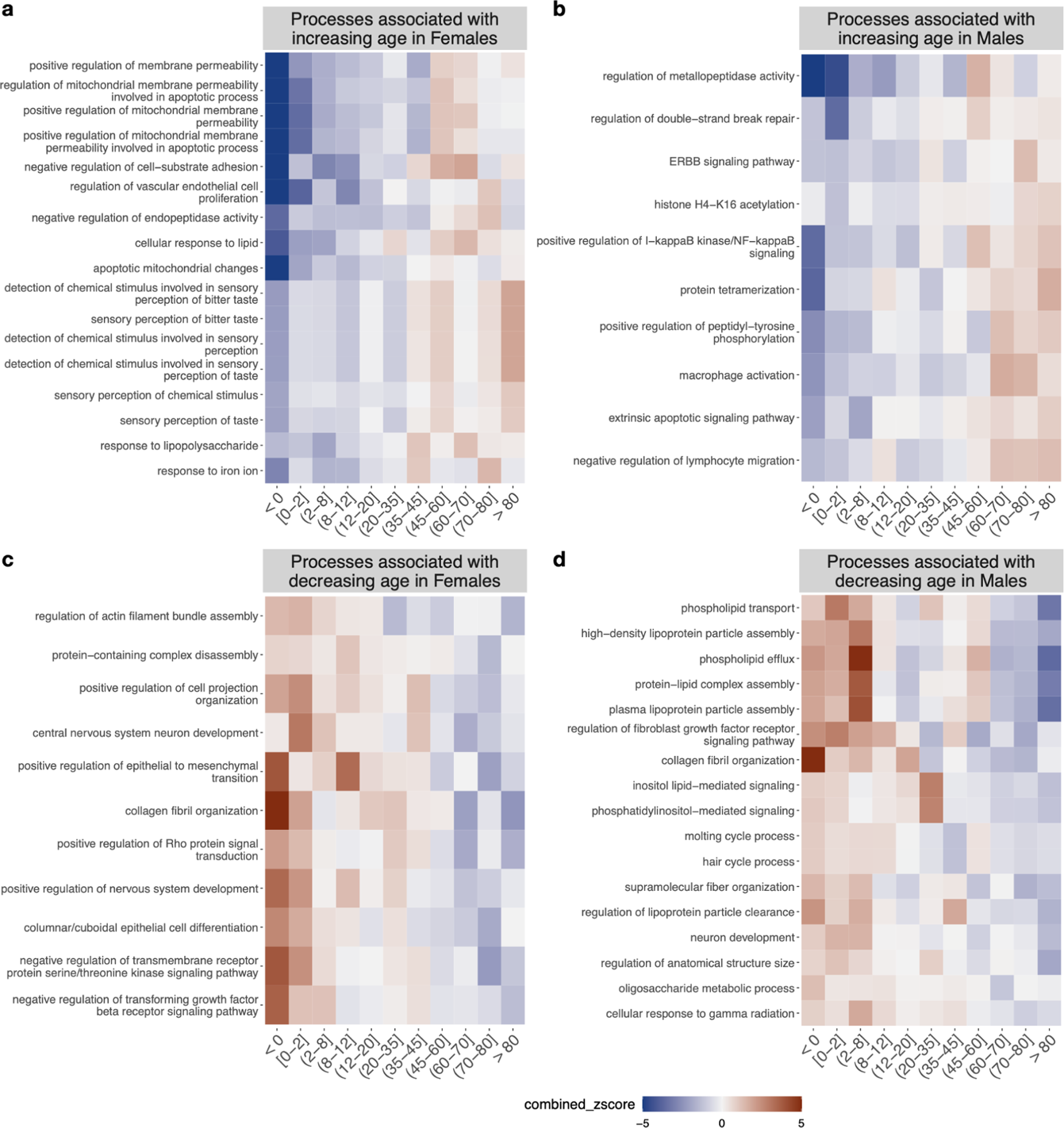
Enrichment of experimentally-derived genes sets from in our sex-stratified age signatures. Age-bias enrichment of experimentally-derived gene sets. Heatmaps show enrichment scores for GO biological processes that are associated with increasing age in (**a**) Females and (**b**) Males and GO biological processes that are associated with decreasing age in (**c**) Females and (**d**) Males. Each of these biological processes has a Spearman correlation over +0.8 or under –0.8 with age group.

On the other hand, processes that are associated with higher expression in younger age groups and lower in older age groups include developmental processes and collagen fibril organization, which has been noted in multiple prior studies [62–64]. Together, these results suggest that our age signatures and enrichment patterns will be powerful for studying sex-specific processes of aging and development. To make these results easily available for biomedical researchers as well as computational researchers, we have created a web-app (http://mlgenesignatures.org/) to display plots and create downloadable tables.

## Discussion

Age and sex have historically not received the attention they deserve in biomedical research, resulting in fundamental gaps in our understanding of how these factors influence normal physiology and disease mechanisms. In this project, we make an effort to enable the biomedical community to systematically address these gaps. First, we assembled the largest set of manually-curated age- and sex-labeled bulk, primary human gene expression samples to date. Using these data, we show that it is possible to predict age-group from gene expression with reasonably high accuracy using simple one-vs-rest logistic regression models. We then investigated the genes across the genome with age-stratified sex bias and sex-stratified age bias — both identified in a data-driven manner. These analyses have provided insights into several aspects of age- and sex-related human gene regulation, cellular pathways, and disease.

### Sex and age group prediction from gene expression

Though we conducted an expansive search for transcriptome samples with sex and age labels, there are several tissue and disease biases in our dataset. As blood is one of the easiest tissues to collect from humans, it is not surprising that the largest set of samples are from blood (**Fig. S2**). Brain, small intestine, liver, lung, and retina all have hundreds of samples each as well. Often, studies separate samples based on tissue and then determine age- and sex-biased genes through differential expression or age prediction. However, even stratifying samples by tissue is not enough to determine age- and sex-biased genes without prejudice as cell type composition affects these results [35,47]. One study that considered multi-tissue signatures in age prediction showed that prediction improves when gene expression from more than one tissue is used to predict age [29]. Ren and Kuan [33] recently compared several different tissue-specific and across-tissue genesets to use as features for expression-based age prediction models using data from GTEx. Their results show that an across-tissue geneset derived from expression across all GTEx tissues have similar performance in prediction to using only differentially expressed genes for a given tissue. When training in one tissue and predicting age in another, the across-tissue feature set was superior. We too tested age group prediction using only blood samples in our dataset and found that prediction performance decreased in most age groups (**Fig. S11-16**). Combined with results from previous studies, our findings suggest that age signal-to-noise ratio improves when including expression from multiple tissues. In determining sex-biased genes, we found that subsetting to only blood samples does not meaningfully change the results (**Fig. S3**). In addition to tissue biases, our dataset also certainly has disease biases due to inherent differences in the incidence of disease in different age and sex groups, along with their likelihood to be studied. The National Institutes of Health lists infectious diseases, brain disorders, and cancer as amongst its top-funded in disease research in recent years [65]. As expected, these diseases make up a large number of the samples in our dataset.

Age group prediction was more difficult in the middle age groups (**Fig. 4b,d**, **Fig. 5b,d, Fig. S5-8**). This is not surprising, as environmental factors, lifestyle, and aging begin to contribute to more heterogeneity in these age groups, although nonlinearly and nonuniformly [66]. This is also reflected in the similarity between age group models (**Fig. S17-21**). There is more similarity between the young age group models than in older age groups, especially when comparing across sexes and technologies. The dissimilarity between RNA-seq and microarray models is likely due to technological differences including the large difference in dynamic range between RNA-seq and microarray experiments. Nevertheless, even if the genes used in the same age group models are different across technologies, it is possible that they play a role in similar biological processes and pathways, which we can test in the future by comparing enrichment results between technologies.

### Enrichment of experimental genesets in age and sex signatures

Our age-stratified sex-signatures showed many immune response-related genesets to be Female-biased (**Fig. 8**). Females usually produce a stronger immune response than Males and this is thought to contribute to their increased susceptibility to autoimmune diseases [57,67]. This increased susceptibility is profound, as Females account for 80% of autoimmune disease occurrence [56,57]. A few autoimmune disease incidence rates are close to even between sexes, but there are no common autoimmune diseases that show a bias towards Male prevalence at the degree that autoimmune diseases like rheumatoid arthritis, lupus and Hashimoto’s show towards Female prevalence [68]. The many immune response-related genesets found to be Female-biased in our signatures might help probe the molecular underpinnings of these differences.

Our age-stratified sex-biased signatures were able to recapitulate previously-observed sex differences in cell type composition. A study of individuals aged 20–35 years found that Females have a higher number and proportion of B cells [58]. Another study conducted by Márquez and colleagues with a range of individuals aged 22–93 years found a Male-specific decline in B cell proportion after the age of 65 [59]. This study also found many lymphoid cells and T cell populations to be more abundant in Females, the latter supporting the result of another study that had found naive T cells specifically to be higher in Females [69]. These cell types are more commonly Female-biased in our age-stratified sex-biased signatures (**Fig. 8d,e**). On the other hand, the previously mentioned Márquez et al study also found myeloid lineage cells (particularly monocytes) to be more abundant in Males [59] (**Fig. 8d,e**). We observe this Male-bias in the younger age groups. Altogether, our signatures show high concordance with sex-biases observed in previous immune studies, suggesting that they will be excellent tools to study other pan-body sex-biased processes and disease mechanisms.

We noted several apoptosis and programmed cell death processes associated with increasing age in our sex-stratified age-biased signatures, which are well known to be associated with the process of aging [60] (**Fig. 9a,b**). Positive regulation of NF-kappaB signaling and negative regulation of lymphocyte migration is associated with increasing age in the Male signatures (**Fig. 9b**). These observations are consistent with studies that link NF-kappaB signaling to the aging process [61] and show that adaptive immune function decreases with age, especially in men [59]. Other processes associated with decreasing age are developmental processes and collagen fibril organization (**Fig. 9c,d**). Several studies have associated increased age with lower collagen levels and lower integrity and increased dysregulation of the collagen network [62–64]. These biologically-meaningful age associations show the utility of these data-driven signatures for investigating molecular processes related to aging and development.

### Availability of data and code

We make the set of ∼30,000 age- and sex-associated curated transcriptome samples and code to reproduce these approaches available via GitHub so that other researchers may build upon them for their own studies. Our genome-wide sex-biased and age-biased gene signatures and the associations of hundreds of biological contexts with these signatures will be searchable by an online webserver to make these results easily accessible for biomedical researchers. The community can use these signatures and associated contexts to explore the age- and sex-biased expression patterns of one or more of their favorite genes, use the sex- and age-biased gene signatures to inform the genes prioritized in new studies, and/or search through the thousands of precalculated enrichment results using the names of pathways, cell types, phenotypes, traits, and diseases of interest to examine the association of constituent genes with sex and age. Finally, we will also make expression values of labeled transcriptomes available via the web interface for biomedical researchers to search and compare gene expression between age and sex groups of interest.

## Methods

### Data collection

We downloaded human microarray gene expression data from the Gene Expression Omnibus (GEO) [15] as raw CEL files. Due to different platforms measuring different genes, we restricted the data to samples from the *Affymetrix Human Genome U133 Plus 2.0 Array*. The CEL files were processed with background subtraction, quantile transformation, and summarization using fRMA [70] based on custom CDF [71] mapping probes to Entrez gene IDs. We downloaded Salmon [72] output files for all human RNA-seq samples available in refine.bio [40] and removed samples with over 50% zero counts. The Salmon-calculated TPM values for the remaining RNA-seq samples were used for all further analysis. We also restricted analysis to genes measured on both platforms, for a total of 18,478 genes.

### Curation of age and sex labels

Age and sex labels were curated for microarray and RNA-seq data with a combination of text mining and manual curation. For the microarray data, we downloaded sample descriptions for our microarray from GEO and used simple text matching to identify samples associated with potential age and sex information. We manually checked these text-matched labels by reading the sample descriptions and verifying that the label was correct and removing erroneously labeled samples. For RNA-seq data, we downloaded metaSRA [73] version 1.8 to identify samples associated with potential age and sex information. We then used ffq [74] to fetch sample accession data from the Sequence Read Archive (SRA) [41] to match the sample identifiers used in metaSRA to the run identifiers used in refine.bio. We manually checked these labels as well by reading sample descriptions obtained from SRA. Both microarray and RNA-seq sample descriptions were also used to remove samples that are not primary human samples, and in the case of RNA-seq, to remove samples that are not bulk samples (i.e. single cell or single nuclei). Examples of removed samples include cell lines, xenografts, and pooled samples.

### Age-stratified sex-biased genes

Sex-biased genes were determined separately in microarray and RNA-seq data. Within each set of data, the expression values of each gene were z-scored across all samples. Samples were divided by age group. For each gene, the z-score value that best separated the male and Female samples within an age group was found by calculating the arithmetic mean of sensitivity and specificity at each value from the minimum to maximum z score in steps of 0.2. Essentially, we considered each z score value as a simple model to predict sex. Every sample with an expression z score above the value was labeled ‘Female’ and every sample with an expression z score below the value was labeled ‘Male’. By reconfiguring the balanced accuracy equation (below) to replace ‘positives’ with ‘Females’ and ‘negatives’ with Males, we create a Female-bias metric for the expression of each gene. The balanced accuracy is the arithmetic mean of sensitivity and specificity of a model and ranges from 0 to 1. In our metric, 1 is the extreme end of Female bias (all Female samples have higher expression than the cutoff value and all male samples have lower expression), 0 is the extreme end of male bias (all male samples have higher expression than the cutoff value and all Female samples have lower expression), and 0.5 is perfectly balanced. The same process was repeated using only samples from blood for the blood-only analysis. For heatmaps of strongly sex-biased genes (**Fig. 2, Fig. S3**), we subtract the Female-bias metric score from 1 to plot the balanced accuracy if Males are considered the more highly-expressed group.

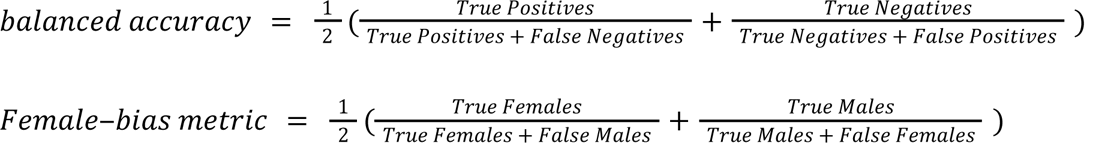

### Logistic regression models

Separately for microarray and RNA-seq data, in each sex, for each age group, we trained a one-vs-rest logistic regression model with an elastic net penalty. RNA-seq data was *asinh* transformed but no other scaling was used. In every sex/technology combination, three folds of all the samples were created for cross-validation by assigning entire datasets at a time, roughly by assigning the three largest remaining datasets to each of the three folds in a manner that kept the number of samples and datasets as equal as possible across all folds. In RNA-seq data, 145 fetal samples were added by predicting sex in samples without sex information to increase the number of samples to a number viable for 3-fold cross validation. Sex was predicted based on 15 genes with over 0.9 balanced accuracy in separating Female and Male samples with a simple expression cutoff. Only samples with sex agreement in at least 13 out of the 15 genes were labeled with the predicted sex and kept in our labeled set.

To create sex-stratified age signatures in blood samples only, we assigned the 3 largest datasets of each age group to one of three test folds making concessions to keep full datasets together if there was a conflict across age groups. To ensure each test fold had at least one dataset with at least 5 samples from a given age group, we excluded age groups without enough data to meet this threshold in both microarray and RNA-seq data. The remaining datasets were used in training for all three test folds. The folds sizes and age group distribution for all models are shown in **Figures S4** and **S10**.

### Curation of other genesets for enrichment analysis

Genesets from previous sex differential expression studies, GTEx, SAGD, and Guo *et al*, were downloaded from the supplemental data in their respective publications. We used all genes declared significant by the authors for any defined group they had curated. Biological Process annotations with experimental evidence codes (EXP, IDA, IPI, IMP, IGI, TAS, IC) were downloaded from the Gene Ontology [49] and propagated to all ancestor Gene Ontology Biological Process (GOBP) terms. We then subset GOBP terms to those with 10 to 200 annotated genes to remove terms with too few genes to do enrichment or terms that are too general to be practically useful. Human disease genes were downloaded from the Monarch Initiative [51,52] at https://data.monarchinitiative.org/latest/tsv/gene_associations/gene_disease.9606.tsv.gz. The genes in the Monarch Initiative file are annotated to disease terms in the Mondo[75] disease ontology. We propagated these annotations to all ancestor Mondo terms and removed any terms without at least 10 genes. Phenotype genes were obtained from Mouse Genome Informatics [76] (MGI) at http://www.informatics.jax.org/downloads/reports/MGI_GenePheno.rpt. We converted these annotations to human genes using the MGI file at http://www.informatics.jax.org/downloads/reports/HOM_MouseHumanSequence.rpt and propagated them to all ancestor terms in the Mammalian Phenotype Ontology [77]. We created sets of genes for GWAS Atlas [50] traits using the Release 3 metadata file at https://atlas.ctglab.nl/#:~:text=Plain%20text%20file%3A-,gwasATLAS_v20191115,-.txt.gz%0AExcel and the MAGMA p value file at https://atlas.ctglab.nl/#:~:text=gwasATLAS_v20191115_magma_P. The genes with the 25 lowest p values were selected as the geneset for each trait. Cell type marker genes from Ianevski et al [54] were downloaded from github at https://github.com/IanevskiAleksandr/sc-type/blob/master/ScTypeDB_full.xlsx.

### Enrichment analysis

In order to determine an enrichment score for each geneset in the age and sex signatures, we used a permutation test to calculate a z score. For each geneset, we calculated the average age-stratified sex enrichment or sex-stratified age bias using the Female-bias metric (converted to a range of –1 to +1 by multiplying it by 2 and subtracting 1) or weight in the logistic regression model, respectively. Then, we pulled a random sample of genes of the same size as the geneset to calculate the average bias of that set. This was repeated 100,000 times and the mean and standard deviation of this distribution was used to calculate a z score for the bias of the original geneset.

The enrichment process for age-stratified sex-biased genes was done separately in the microarray and RNA-seq data using their respective converted Female-bias metric scores for all genes. The z score obtained for each geneset in microarray and RNA-seq data is combined via Stouffer’s method (equation below). For sex-stratified age enrichment, the weight of each gene in the one-vs-rest logistic regression models were used in the enrichment process. As there were six trained models for each age group (3 RNA-seq models, 3 microarray models), we used our permutation test to determine a z score for each experimentally-derived geneset for each model, averaged the z scores across the 3 folds in each technology separately, and combined the z score from RNA-seq and microarray with Stouffer’s Method. This final z score was taken as the enrichment score for each age group (equation below).

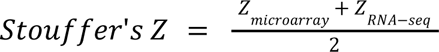

For visualization, extreme z scores were reduced to +5 or –5 if they were higher or lower than that, respectively. The set of representative genesets/ontology terms for visualization were generated using *orsum* [78], which, given the full list of genesets discards any geneset/term if there is a more significant term that annotates at least the same genes.

## Availability of data and materials

The expression dataset identifiers and age/sex labels used in this study, along with code to reproduce the analysis, are available at https://github.com/krishnanlab/Age-sex_signatures_in_humans_code. The actual expression data can be downloaded from GEO and refine.bio, as outlined in *Data Collection* in the *Methods* section or accessed at this Zenodo record: https://zenodo.org/doi/10.5281/zenodo.10056217. The expression datasets, gene signatures, and enrichment results can be easily retrieved and visualized using our web-app at http://mlgenesignatures.org/.

## Declarations

### Competing interests

The authors declare that they have no competing interests.

### Funding

This work was primarily supported by US National Institutes of Health (NIH) grants R35 GM128765 to A.K.

### Author contributions

KAJ and AK designed the study. KAJ performed all the analyses. KAJ and AK interpreted the results and wrote the final manuscript.

## Acknowledgements

We thank the members of the Krishnan Lab for helpful discussion and Alex McKim in particular for sharing the code for performing the enrichment analysis. We are also grateful to the hundreds of labs that contributed microarray and RNA-seq samples to public repositories, allowing for their reuse in this work.

## Notes

https://github.com/krishnanlab/Age-sex_signatures_in_humans_code

https://zenodo.org/doi/10.5281/zenodo.10056217

http://mlgenesignatures.org/

### Competing Interest Statement

The authors have declared no competing interest.

### Summary of Updates

Added link to a new web-app that researchers can use to retrieve, explore, and download our results.

