## Supplementary Figures and Tables for "Human pan-body age- and sex-specific molecular phenomena inferred from public transcriptome data using machine learning"

Supplemental materials

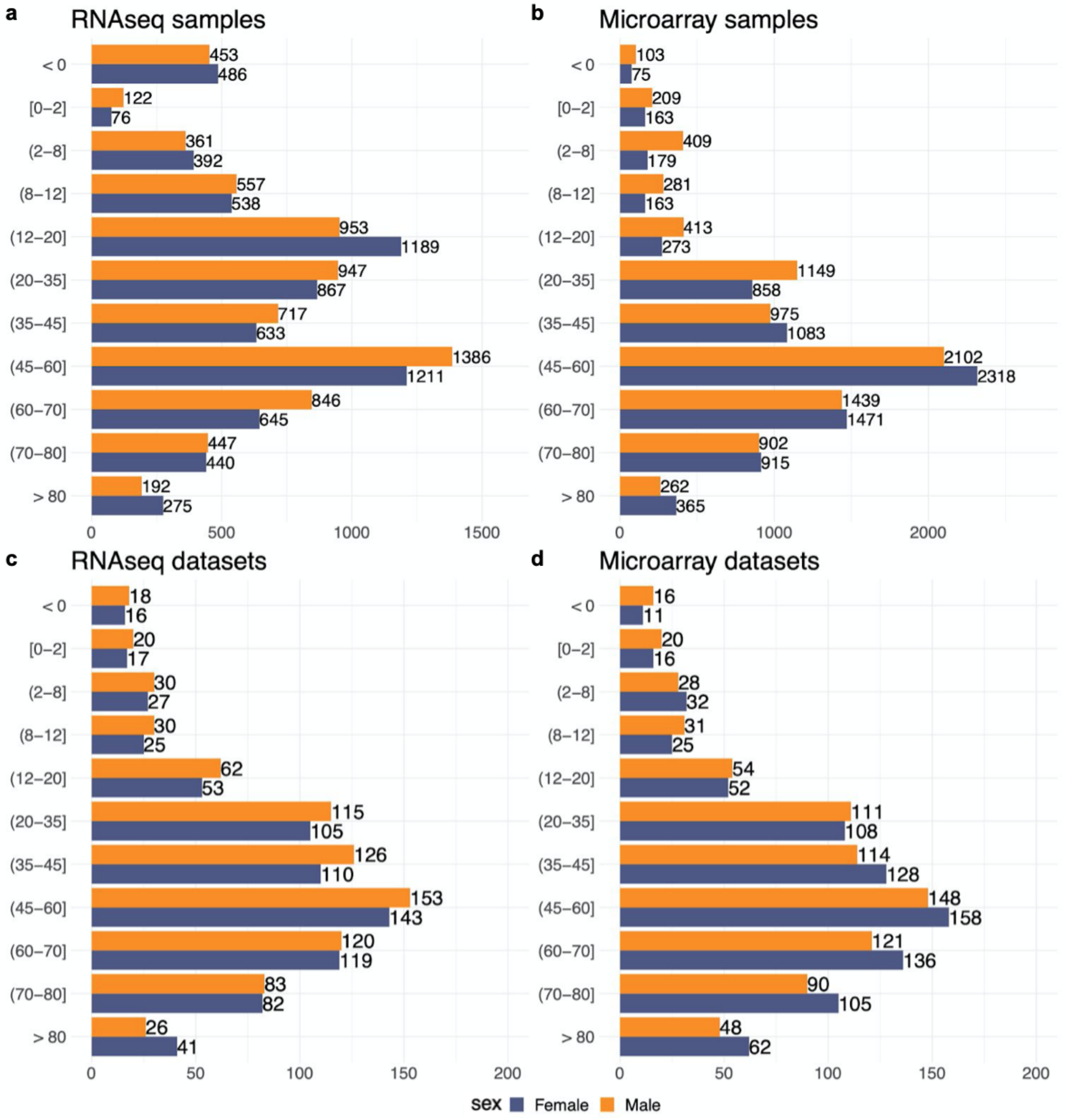


**Figure S1. Number of samples and datasets across age and sex groups.** Number of samples labeled with age and sex in (**a**) RNAseq and (**b**) microarray. Number of datasets in (**c**) RNAseq and (**d**) microarray.

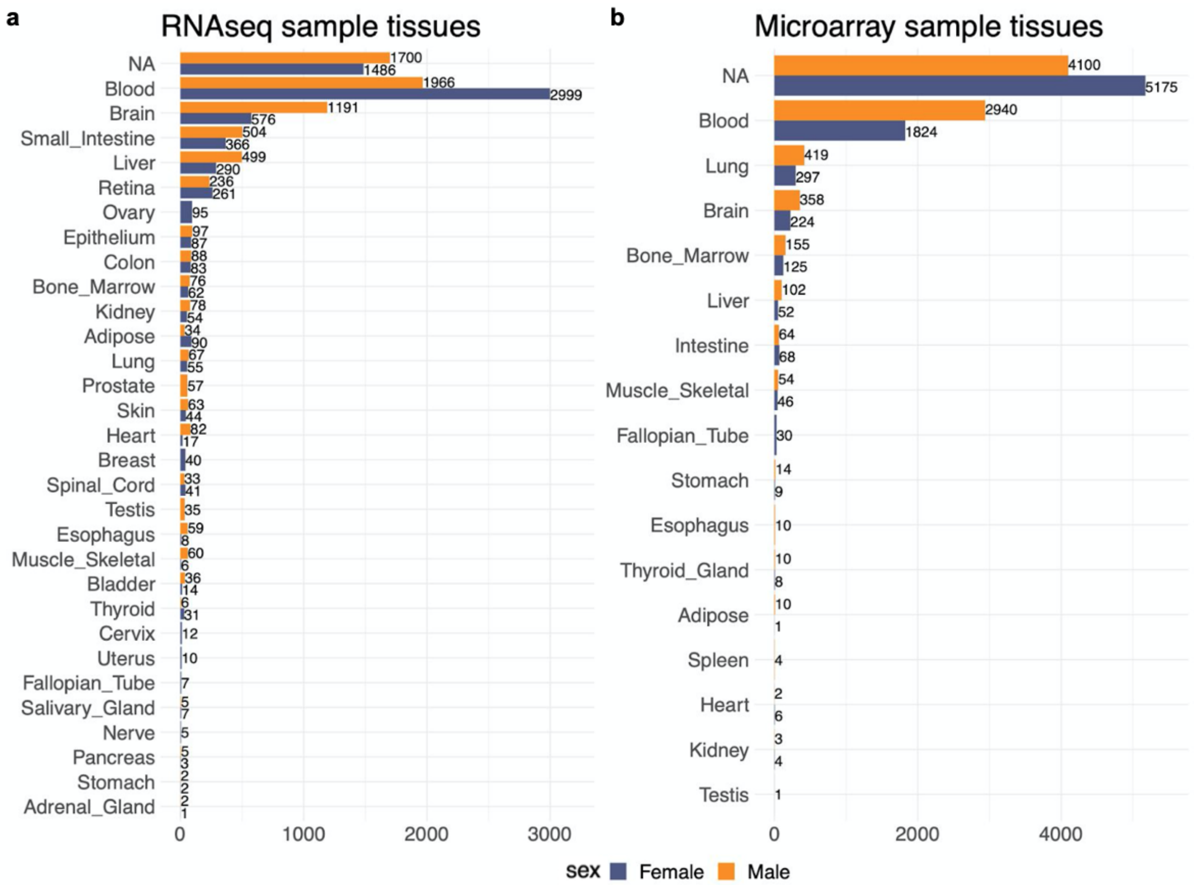


**Figure S2. Number of samples across tissues.** Number of samples per tissue in (**a**) RNAseq and (**b**) microarray.

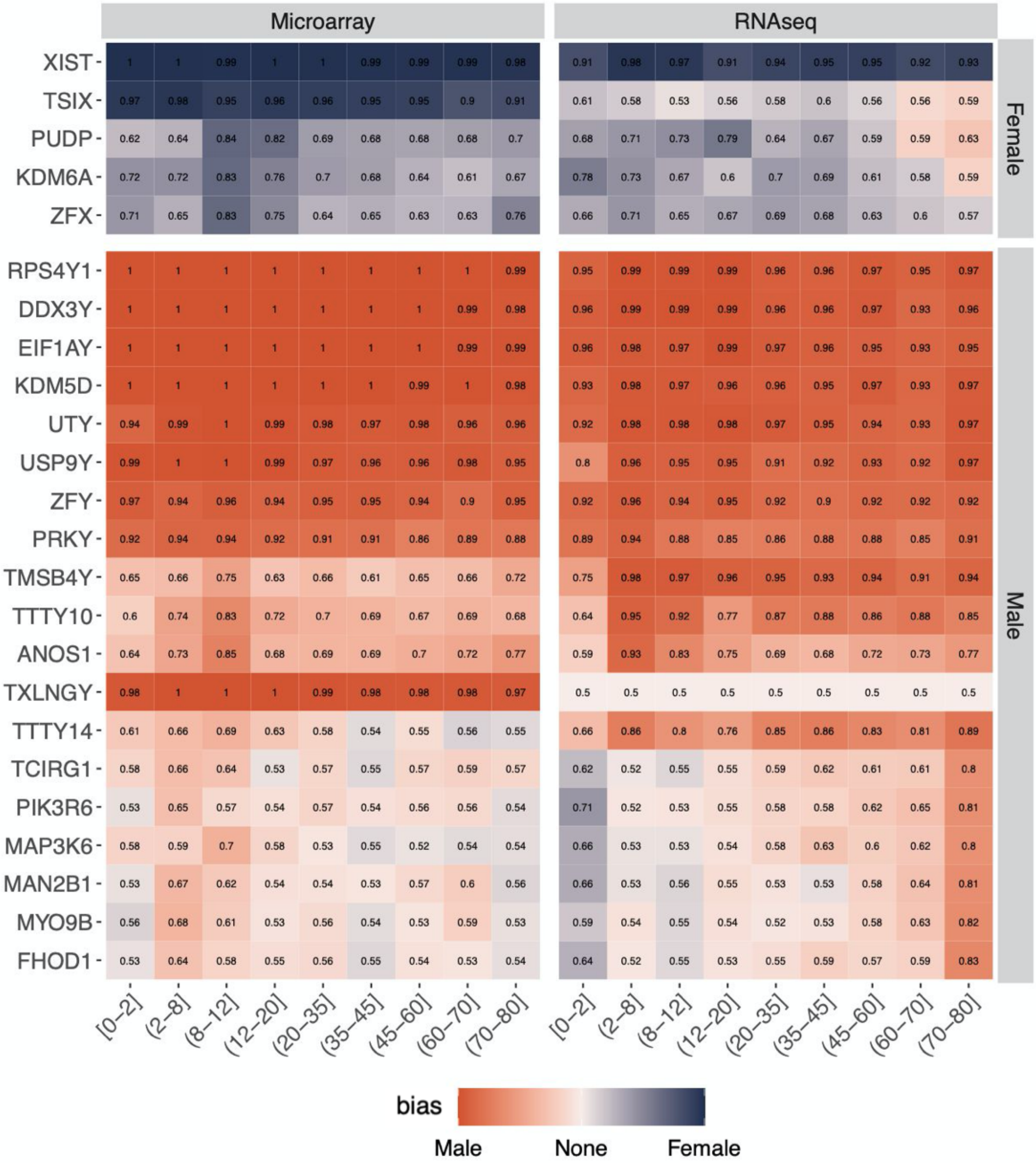


**Figure S3. Most strongly sex-biased genes in blood samples across age ranges.** The heatmap displays all genes (x axis) that had a balanced accuracy of at least 0.8 in any age range (y axis) when separating Female and Male microarray or RNAseq blood samples.

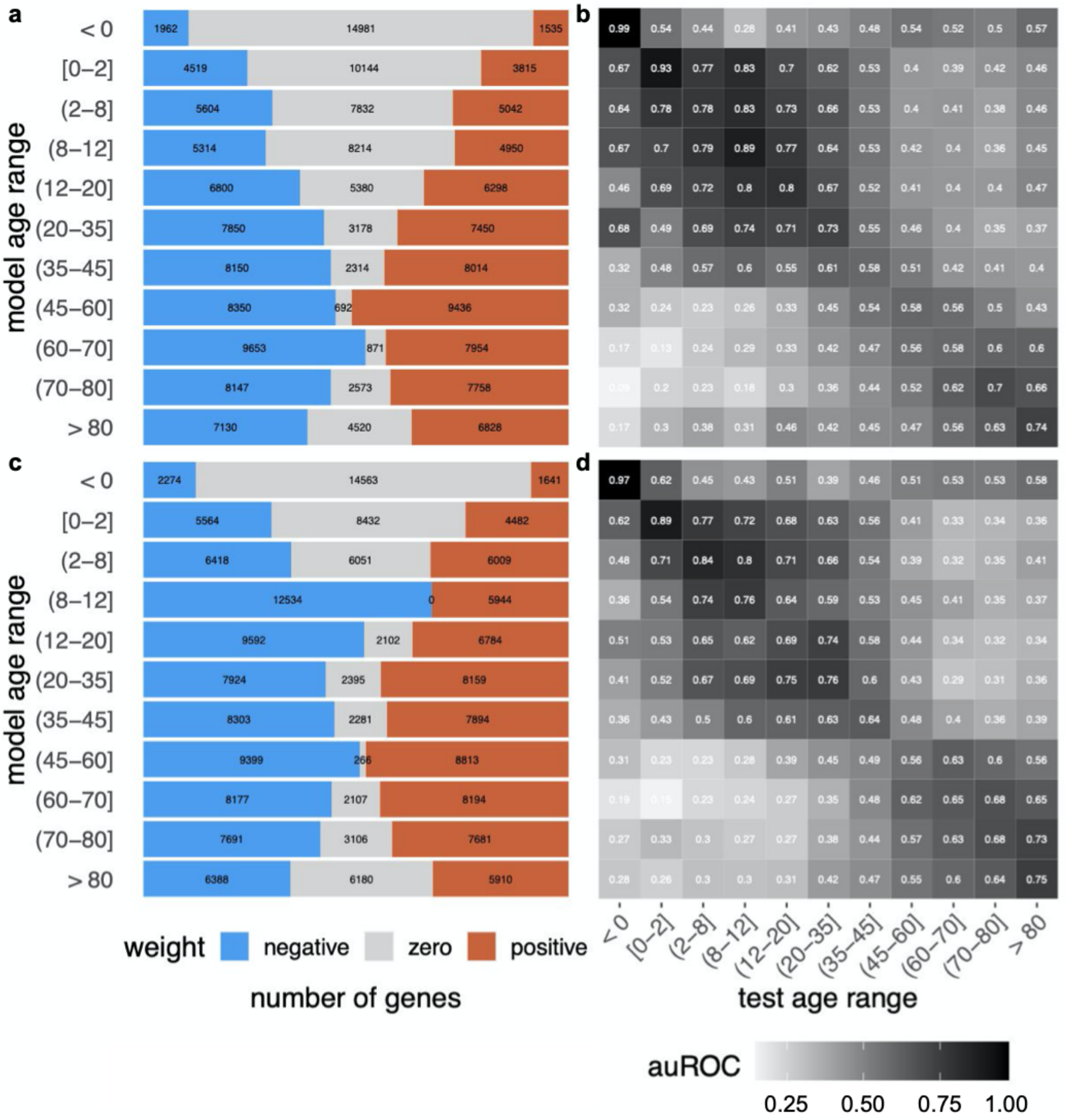


**Figure S4. Size and performance of microarray age group prediction models.** The stacked barplots show the distribution of positive, zero, and negative weights for the model with the median number of positives across the three folds for each age group in microarray for (**a**) Females and (**c**) Males. The heatmaps contain the average auROC of microarray models trained on the age group labeled in the rows when evaluated using the samples in the age group labeled in the column as positive examples for (**b**) Females and (**d**) Males.

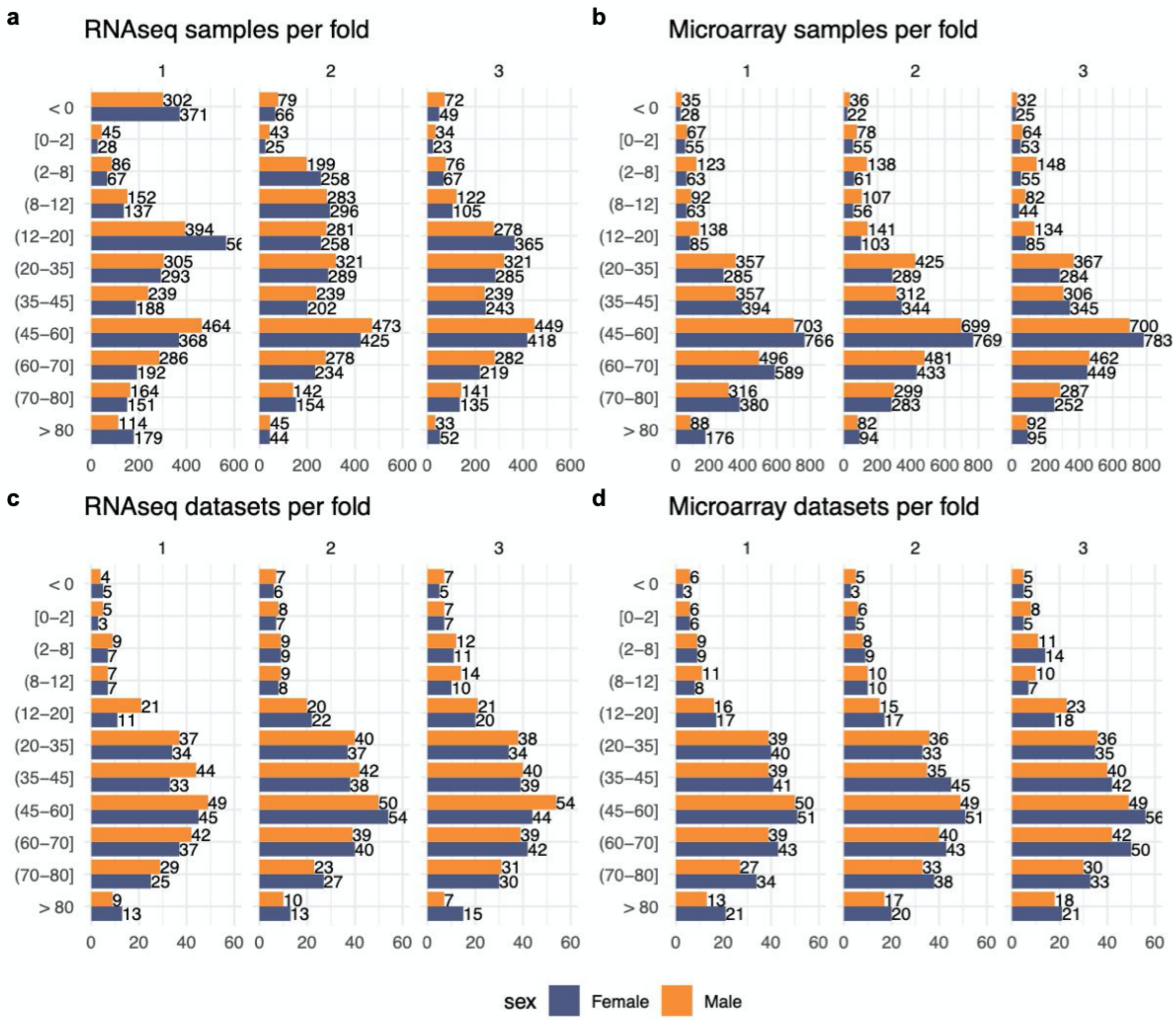


**Figure S5. Fold sizes for age range prediction models.** The top barplots show the number of samples in each fold for models trained in (**a**) RNAseq and (**b**) microarray. The bottom barplots show the number of datasets in each fold for models trained in (**c**) RNAseq and (**d**) microarray.

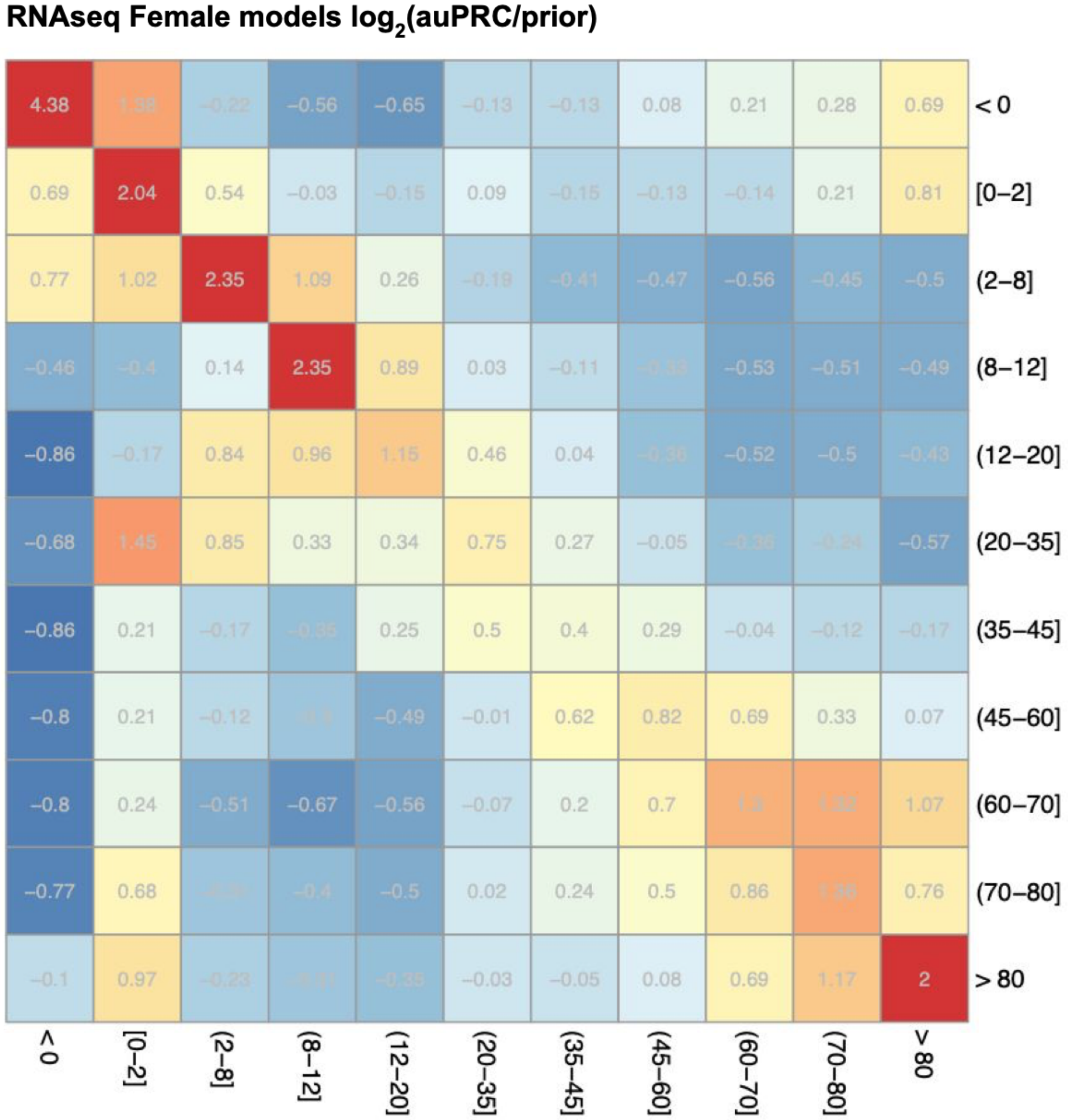


**Figure S6. Performance of RNAseq Female age range prediction models.** The heatmaps contain average $log_2$(auPRC/prior) performance of RNAseq models across 3 folds trained in Female samples on the age range labeled in the rows while evaluated as if the age range labeled in the column were the positive examples.

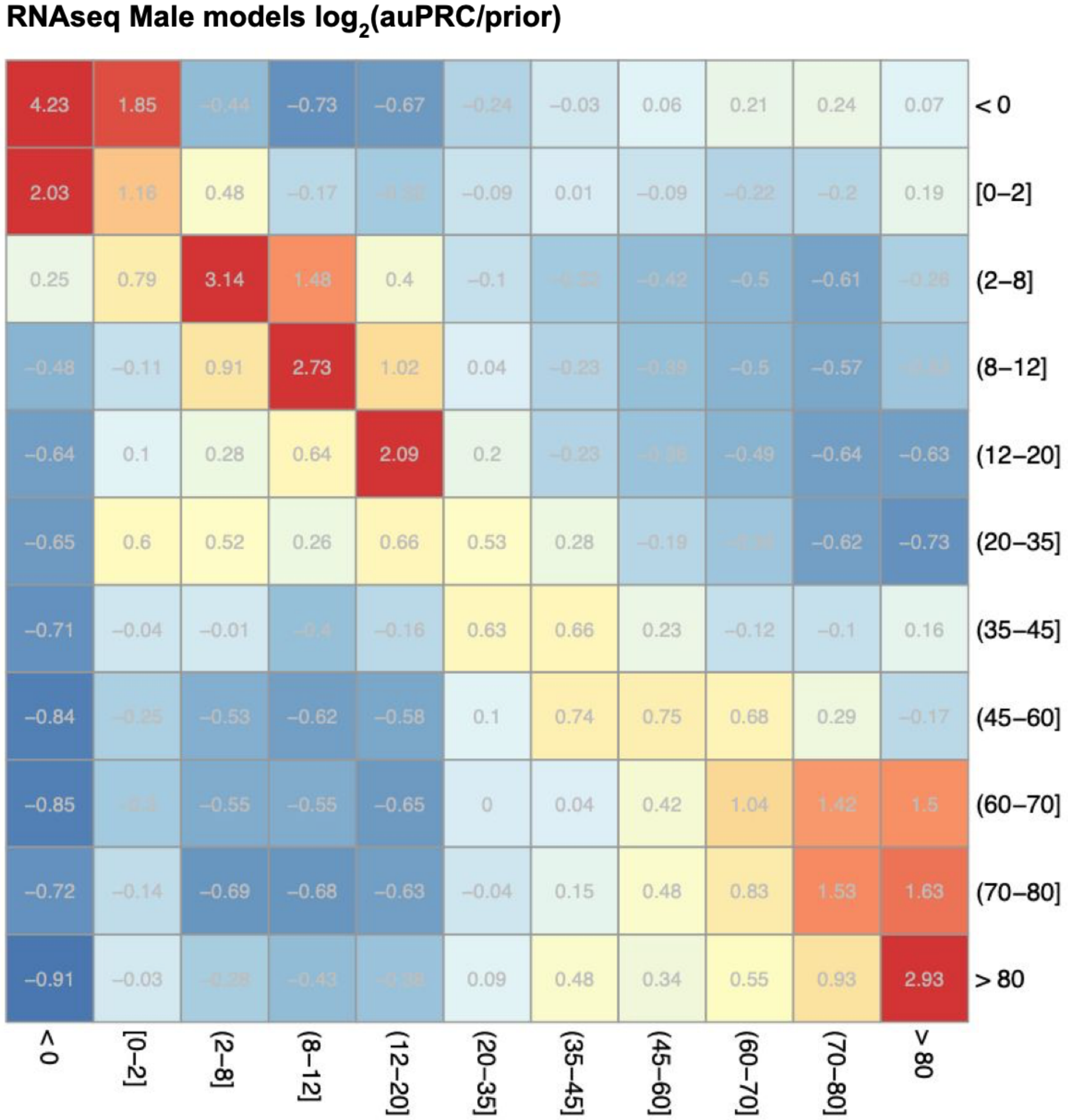


**Figure S7. Performance of RNAseq Male age range prediction models.** The heatmaps contain average $\log_2$(auPRC/prior) performance of RNAseq models across 3 folds trained in Male samples on the age range labeled in the rows while evaluated as if the age range labeled in the column were the positive examples.

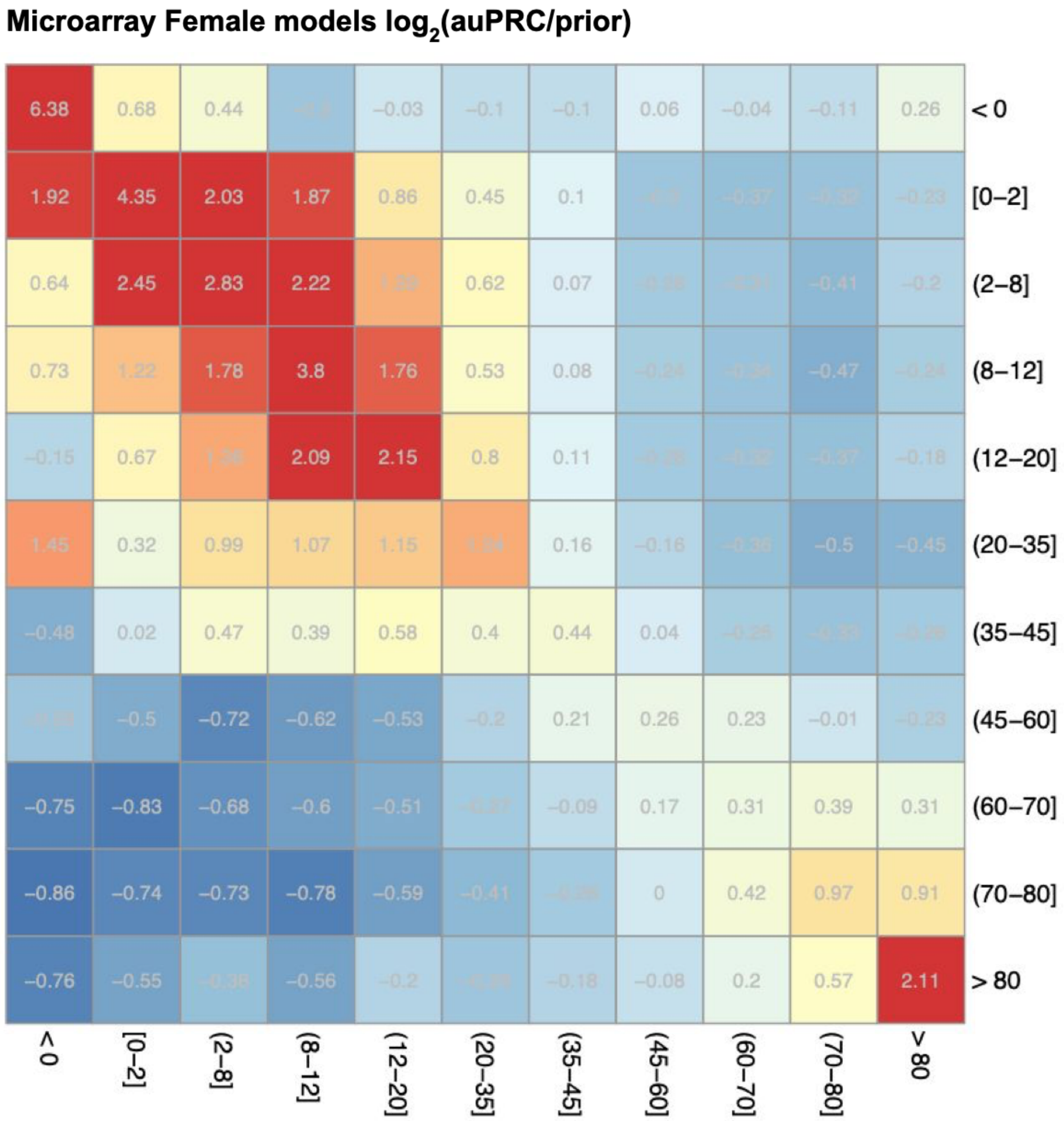


**Figure S8. Performance of microarray Female age range prediction models.** The heatmaps contain average $\log_2$(auPRC/prior) performance of microarray models across 3 folds trained in Female samples on the age range labeled in the rows while evaluated as if the age range labeled in the column were the positive examples.

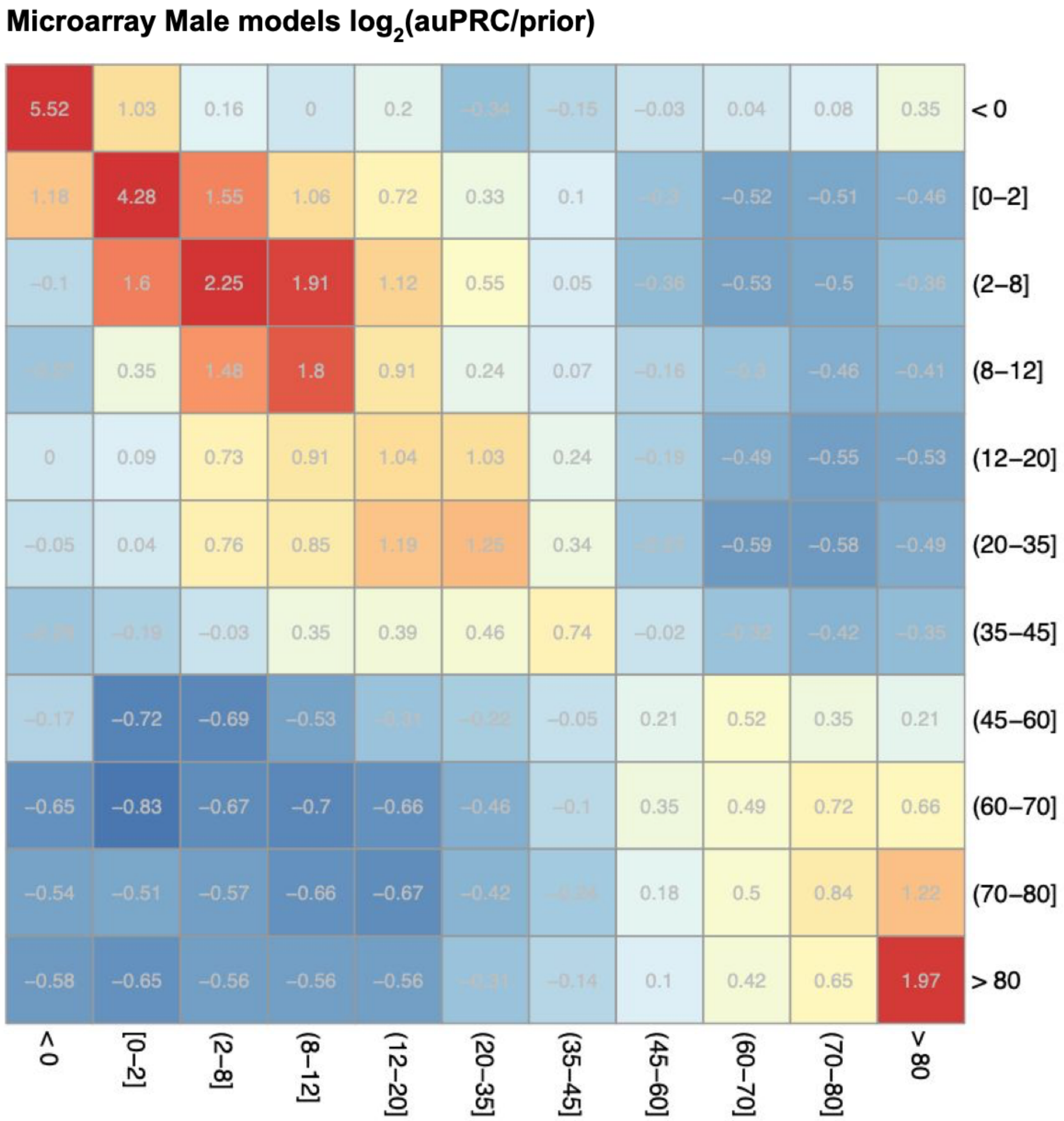


**Figure S9. Performance of microarray Male age range prediction models.** The heatmaps contain average $log_2$(auPRC/prior) performance of microarray models across 3 folds trained in Male samples on the age range labeled in the rows while evaluated as if the age range labeled in the column were the positive examples.

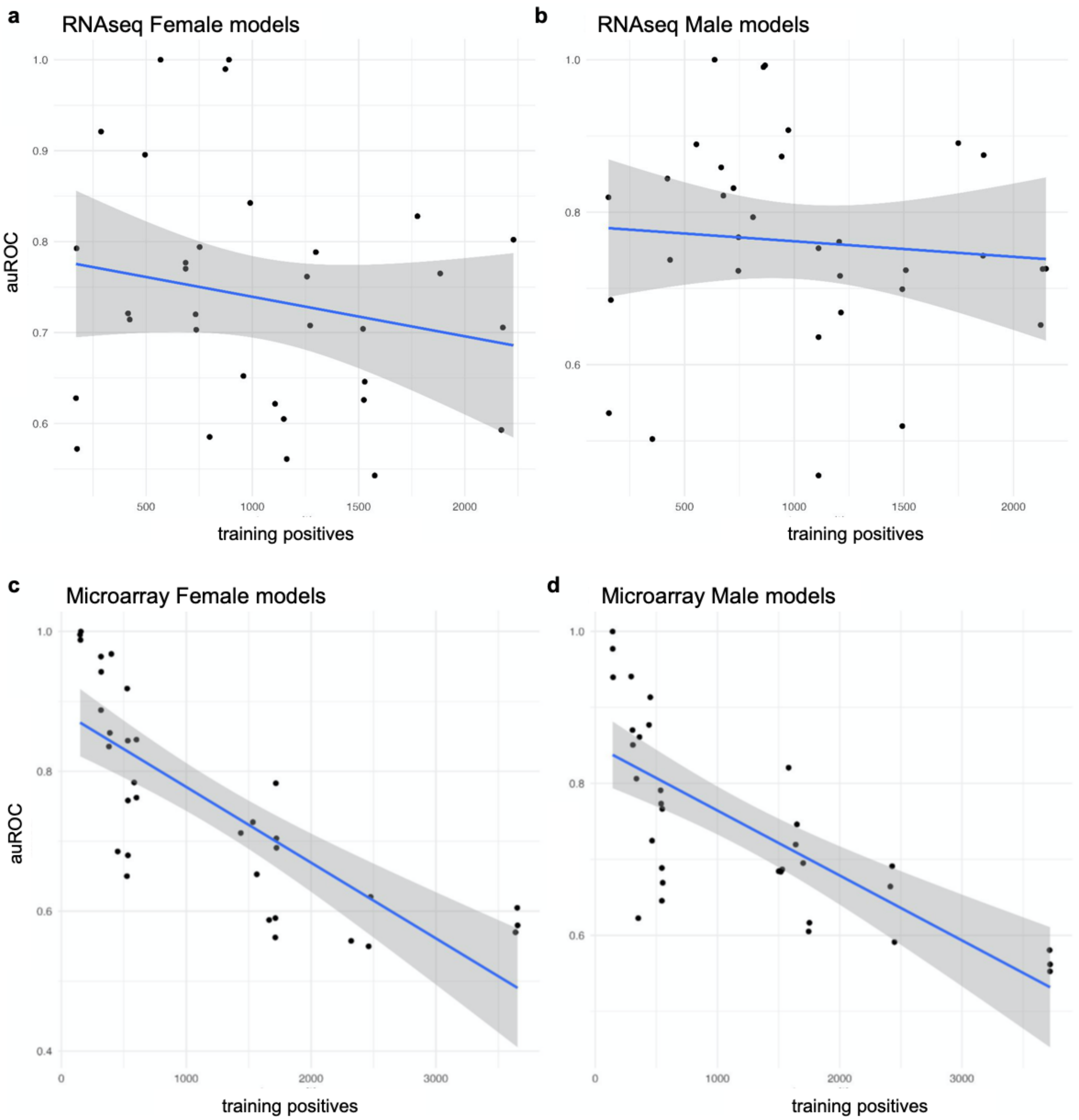


**Figure S10. Number of positive training examples vs auROC performance of age group models.** Scatterplot of the number of positive training examples vs the auROC performance of all RNAseq and microarray Female and Male models.

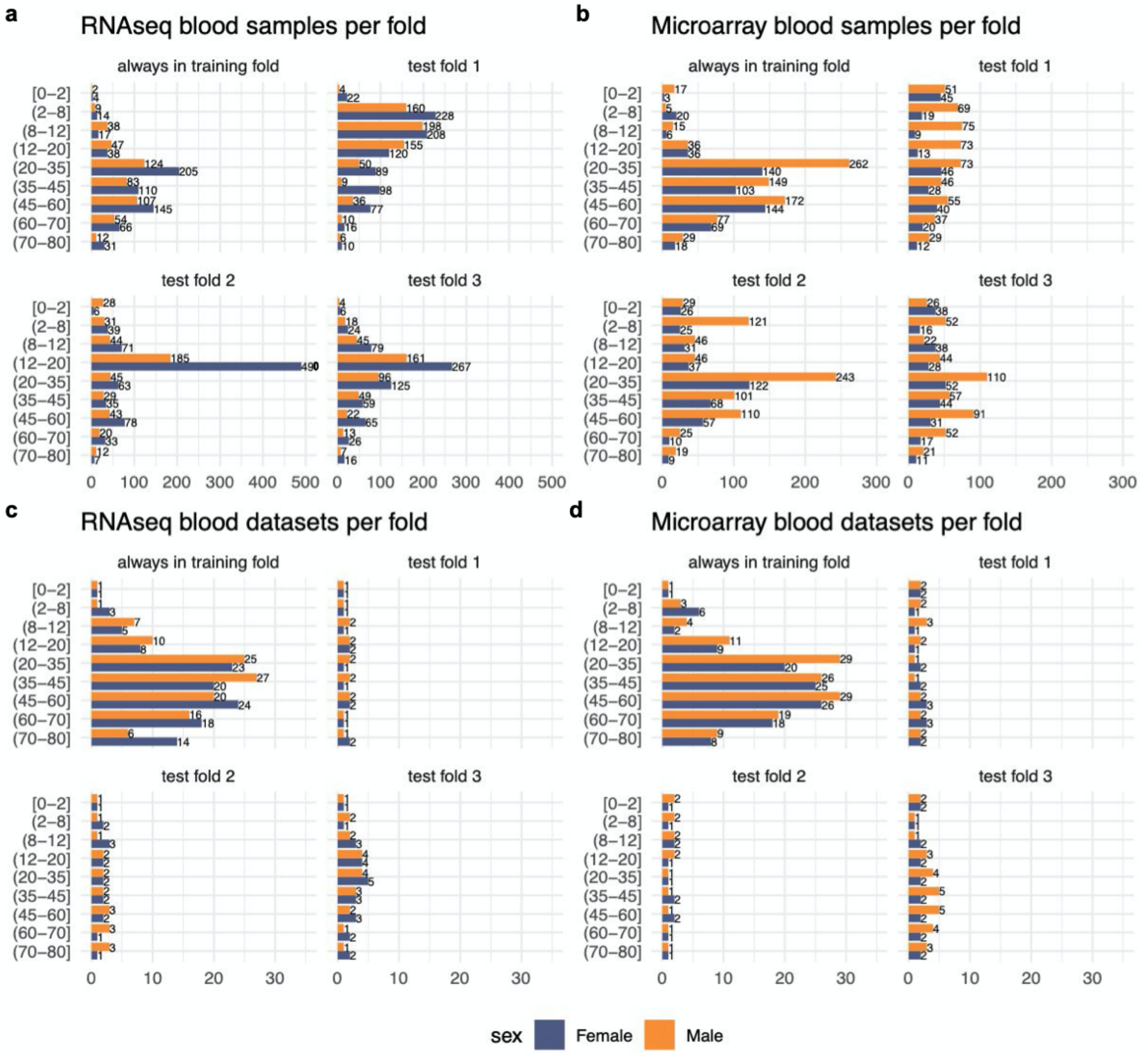


**Figure S11. Fold sizes for age range prediction models in blood samples.** The top barplots show the number of samples in each fold for models trained on blood samples in (**a**) RNAseq and (**b**) microarray. The bottom barplots show the number of datasets in each fold for models trained on blood samples in (**c**) RNAseq and (**d**) microarray.

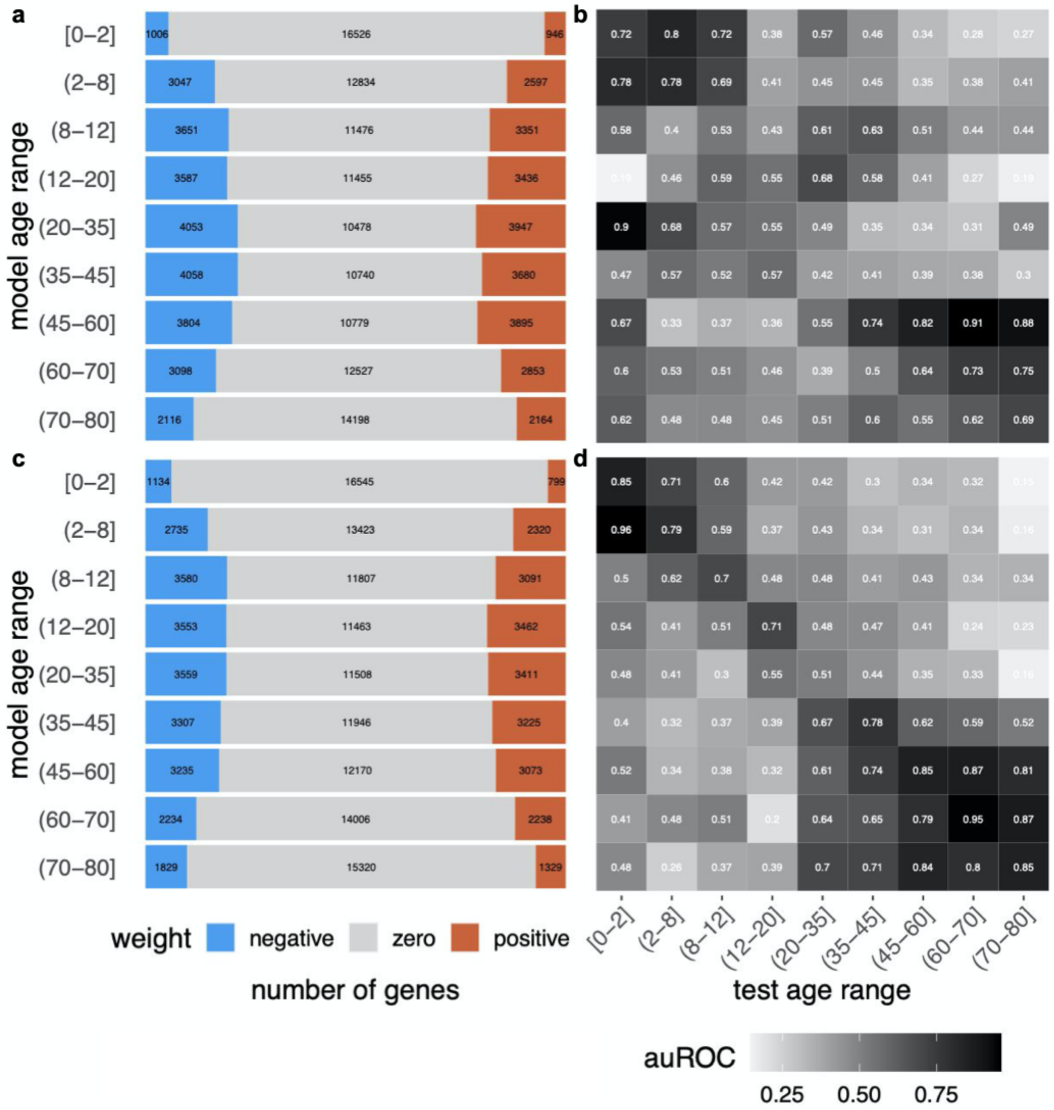


**Figure S12. Size and performance of RNA-seq age range prediction models in blood samples.** The stacked barplots show the distribution of positive, zero, and negative weights for the model with the median number of positives across the three folds for each age range in RNAseq for blood samples from (**a**) Females and (**b**) Males. The heatmaps contain auROC performance of RNAseq models trained on the age range labeled in the rows while evaluated as if the age range labeled in the column were the positive examples (**c**) Females and (**d**) Males.

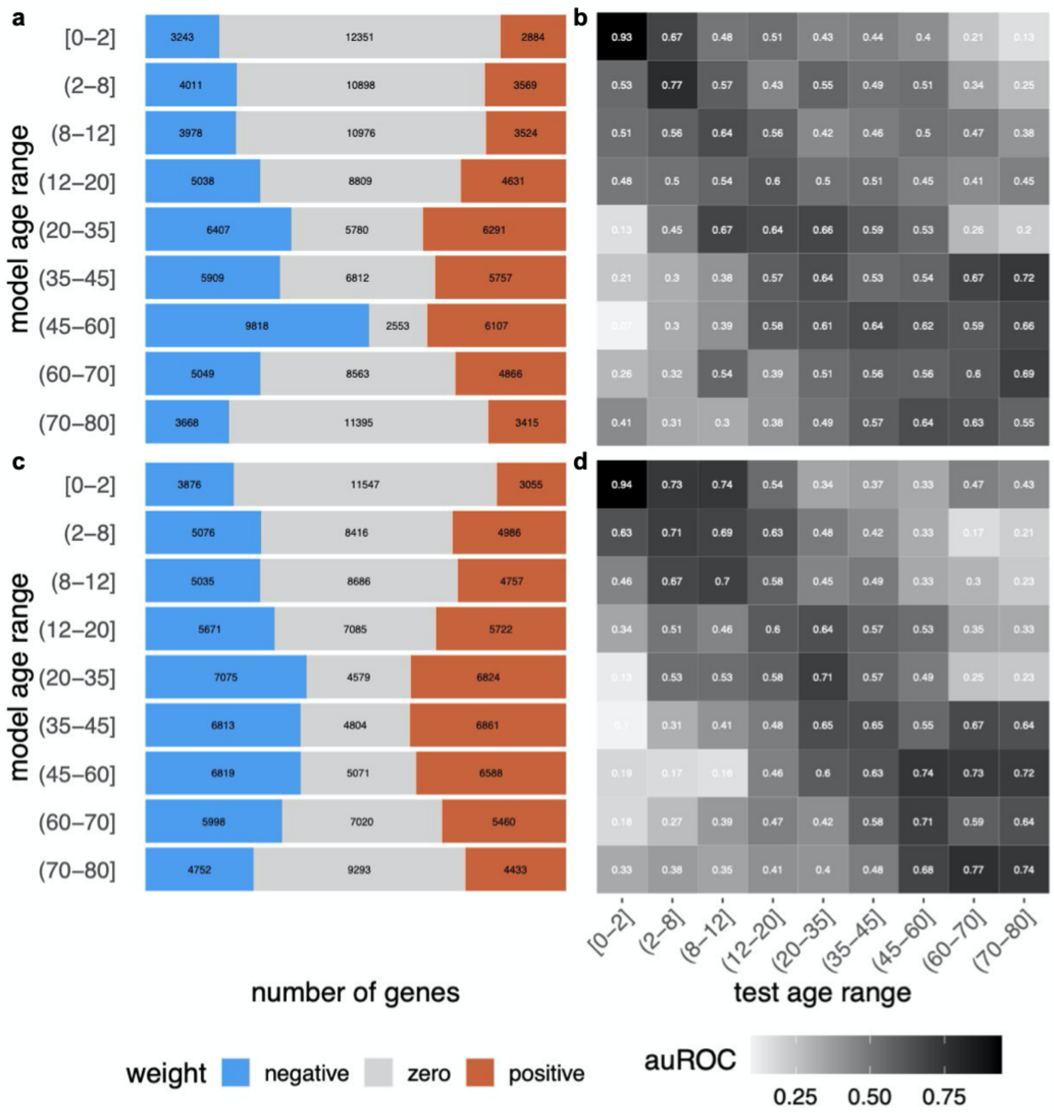


**Figure S13. Size and performance of microarray age range prediction models in blood samples.** The stacked barplots show the distribution of positive, zero, and negative weights for the model with the median number of positives across the three folds for each age range in microarray for blood samples from (**a**) Females and (**b**) Males. The heatmaps contain auROC performance of microarray models trained on the age range labeled in the rows while evaluated as if the age range labeled in the column were the positive examples (**c**) Females and (**d**) Males.

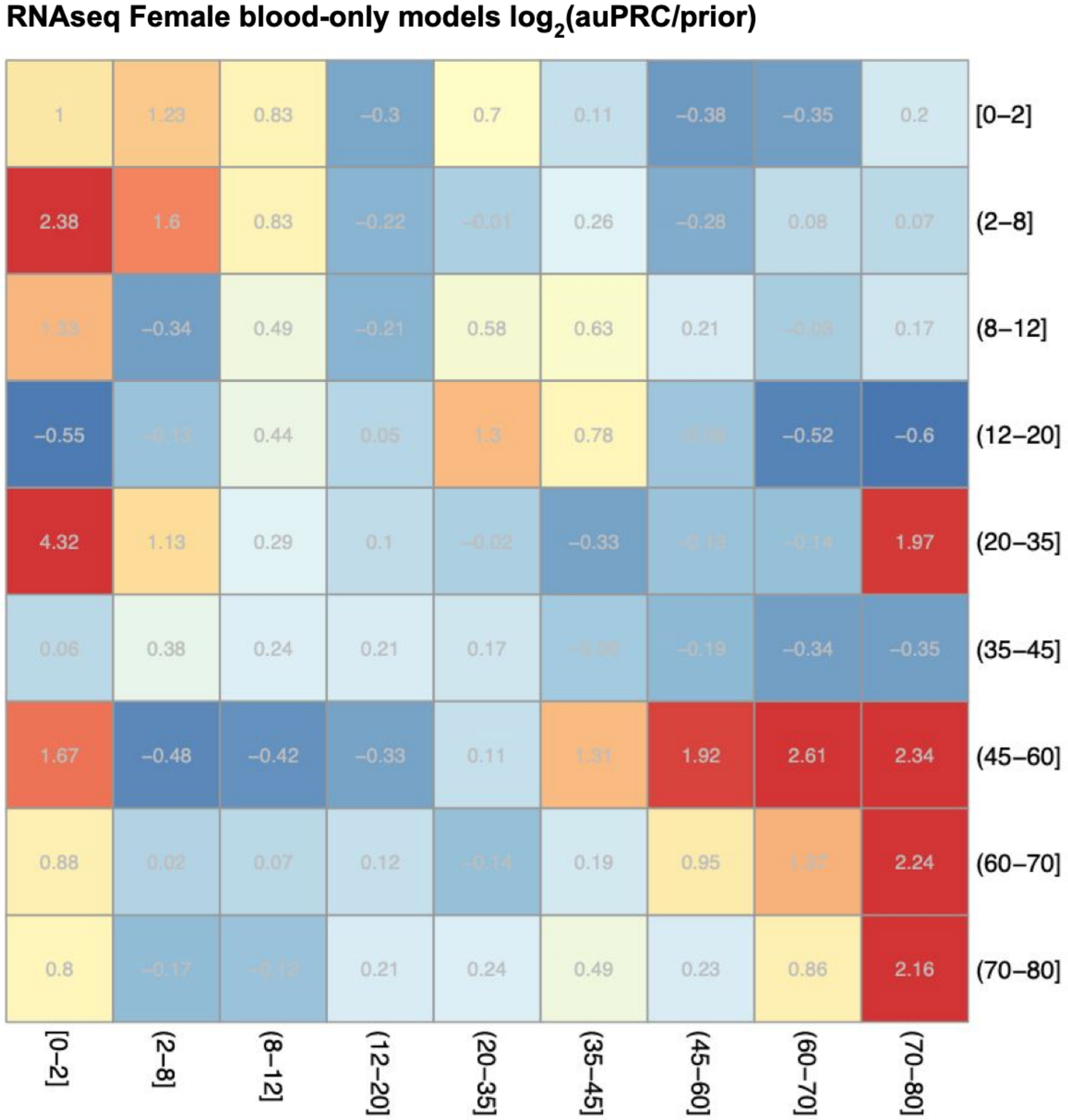


**Figure S14. Performance of RNAseq Female age range prediction models for blood samples.** The heatmaps contain average $\log_2$(auPRC/prior) performance of RNAseq models across 3 folds trained in Female blood samples on the age range labeled in the rows while evaluated as if the age range labeled in the column were the positive examples.

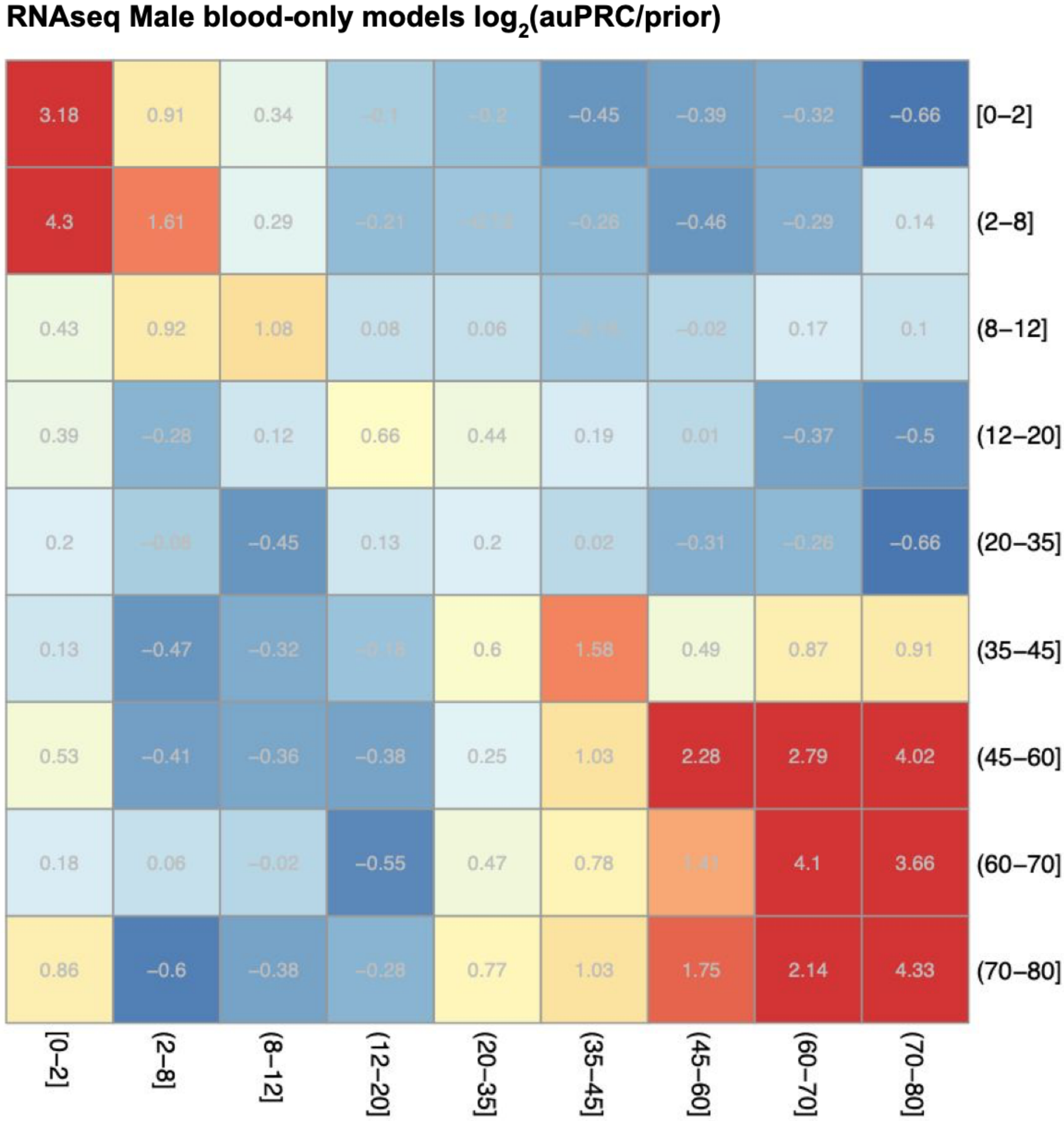


**Figure S15. Performance of RNAseq Male age range prediction models for blood samples.** The heatmaps contain average $\log_2$(auPRC/prior) performance of RNAseq models across 3 folds trained in Male blood samples on the age range labeled in the rows while evaluated as if the age range labeled in the column were the positive examples.

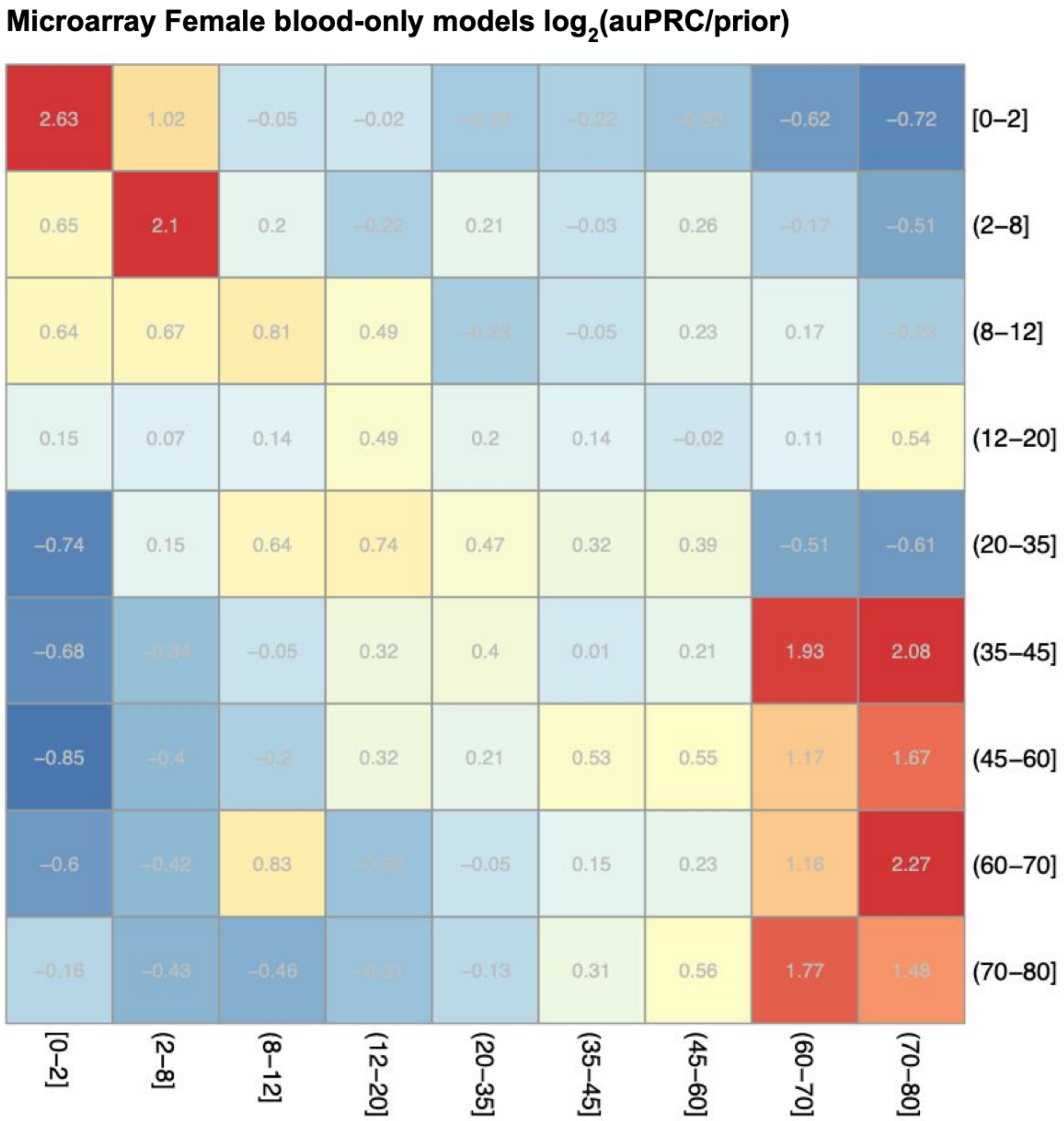


**Figure S16. Performance of microarray Female age range prediction models for blood samples.** The heatmaps contain average $\log_2$(auPRC/prior) performance of microarray models across 3 folds trained in Female blood samples on the age range labeled in the rows while evaluated as if the age range labeled in the column were the positive examples.

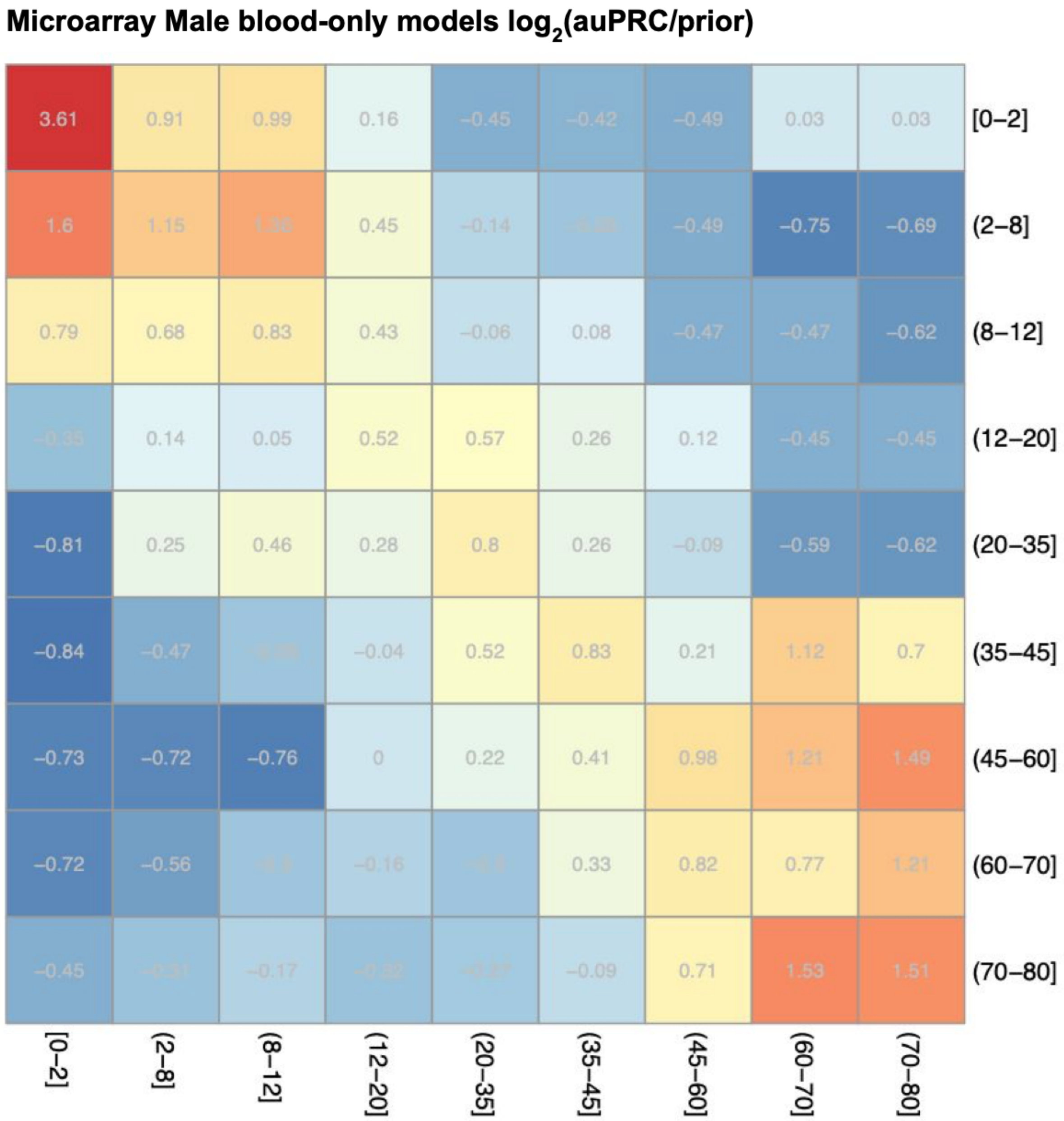


**Figure S17. Performance of microarray Male age range prediction models for blood samples.** The heatmaps contain average $\log_2$(auPRC/prior) performance of microarray models across 3 folds trained in Male blood samples on the age range labeled in the rows while evaluated as if the age range labeled in the column were the positive examples.

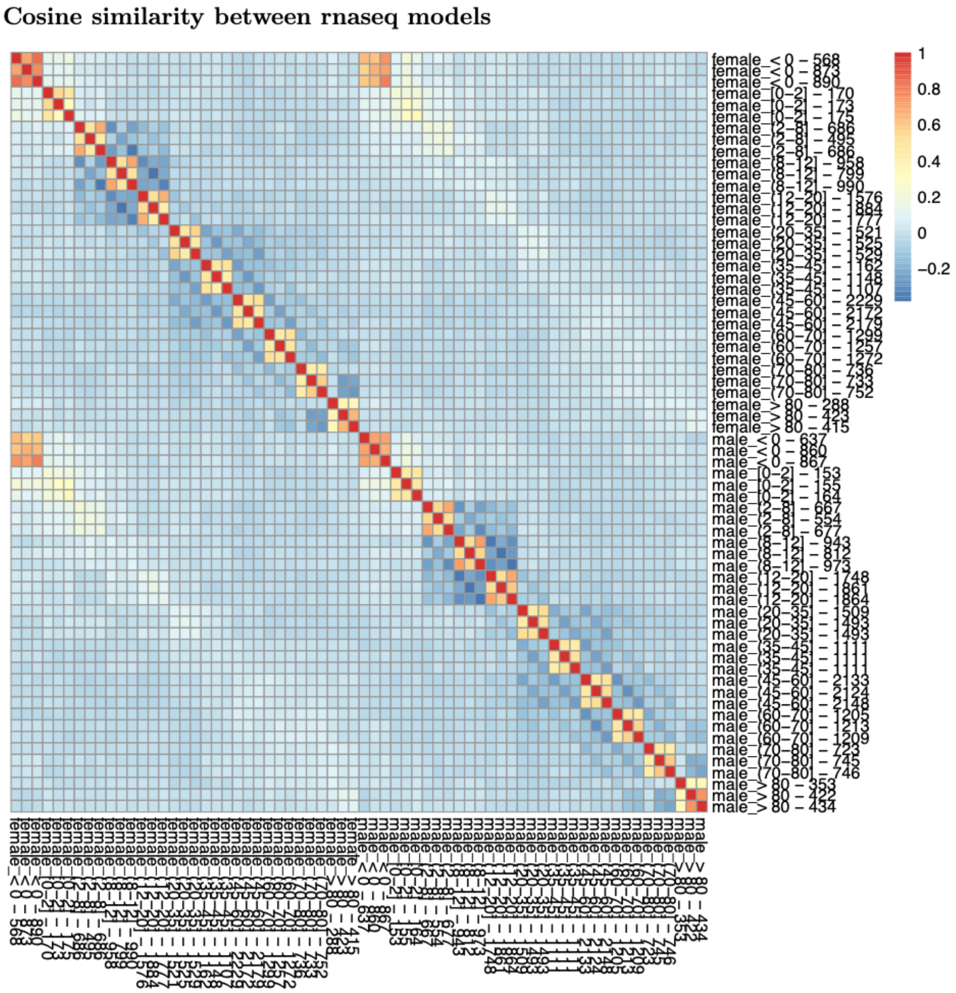


**Figure S18. Cosine similarity of RNAseq model weights.** The heatmap shows the similarity between all RNAseq models trained in both Females and Males. The number after the age range is the number of positive training examples.

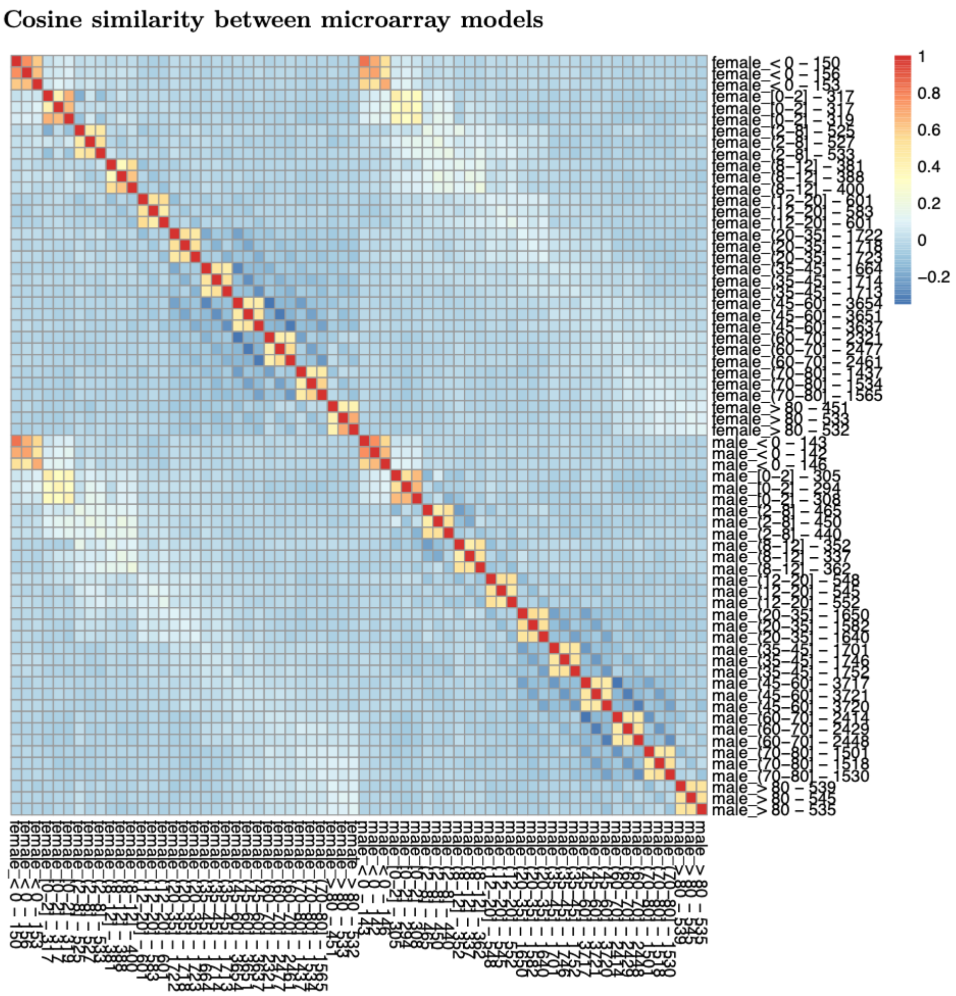


**Figure S19. Cosine similarity of microarray model weights.** The heatmap shows the similarity between all microarray models trained in both Females and Males. The number after the age range is the number of positive training examples.

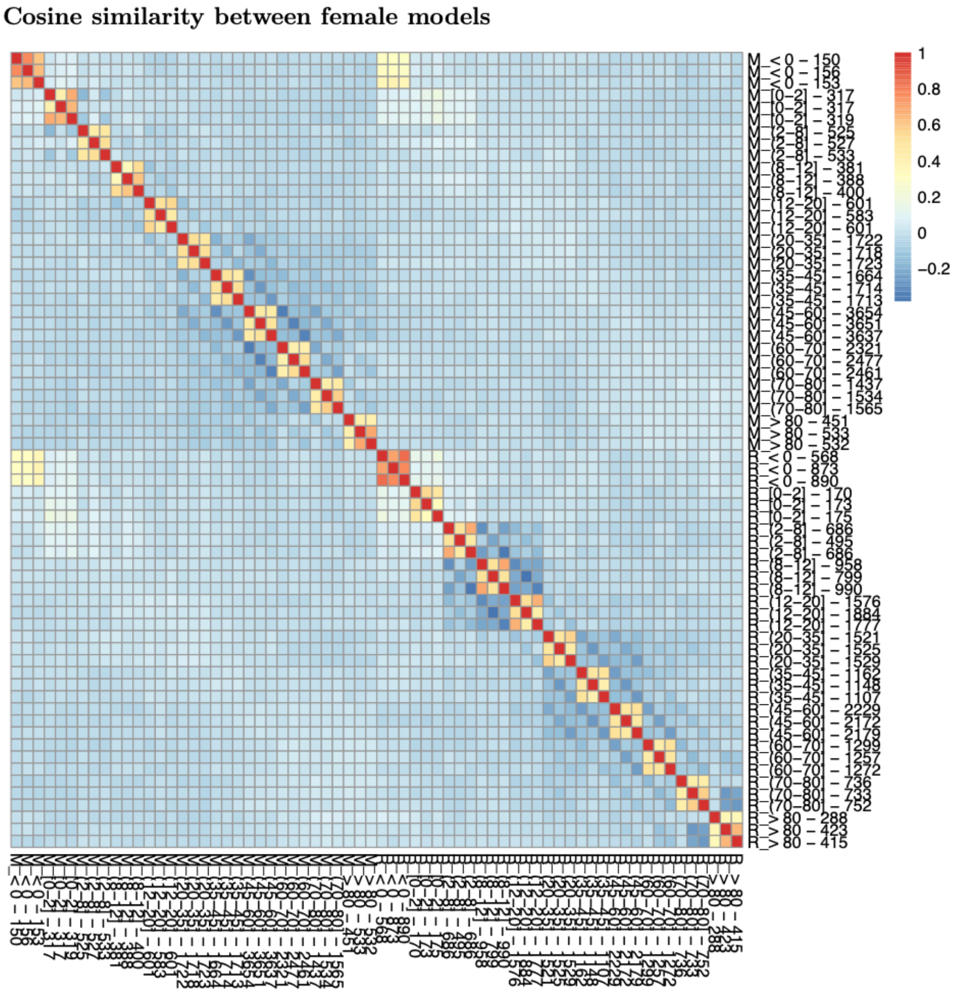


**Figure S20. Cosine similarity of Female model weights.** The heatmap shows the similarity between all RNAseq ('R_' prefix) and microarray ('M_' prefix) models trained in Females. The number after the age range is the number of positive training examples.

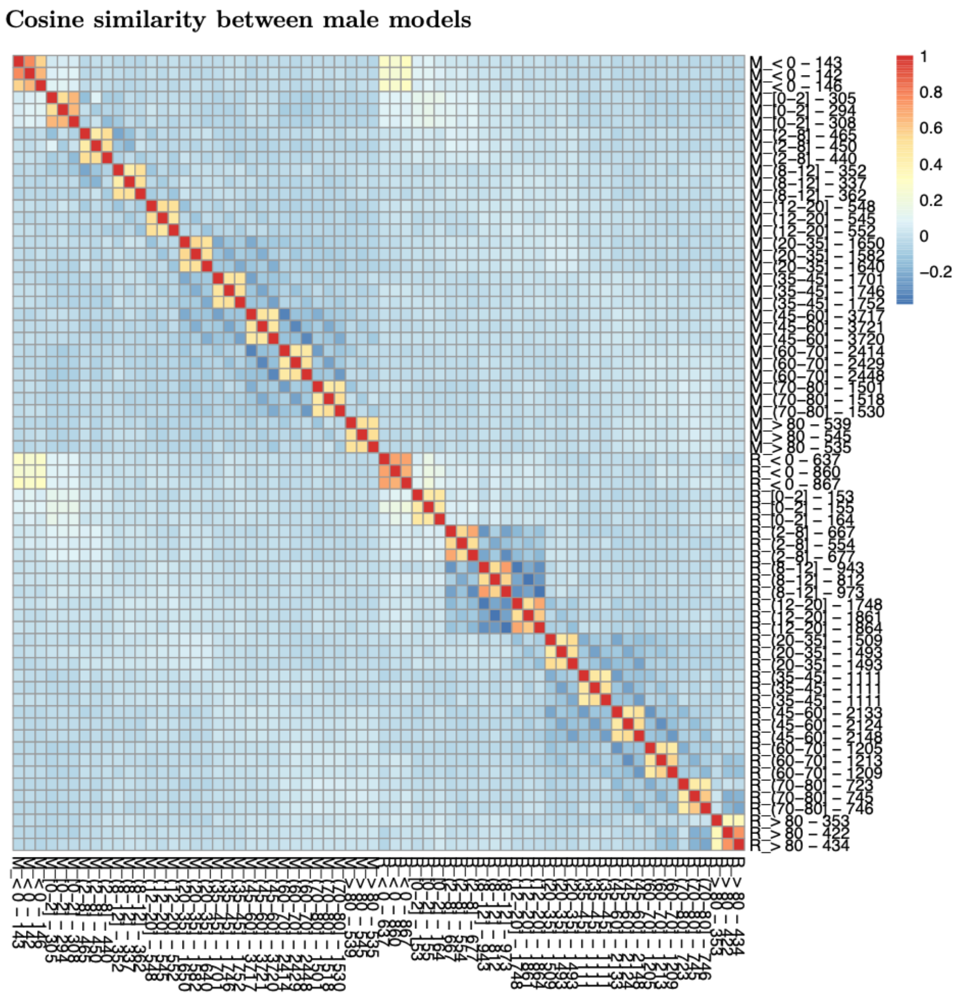


**Figure S21. Cosine similarity of Male model weights.** The heatmap shows the similarity between all RNAseq (‘R_’ prefix) and microarray (‘M_’ prefix) models trained in Males. The number after the age range is the number of positive training examples.

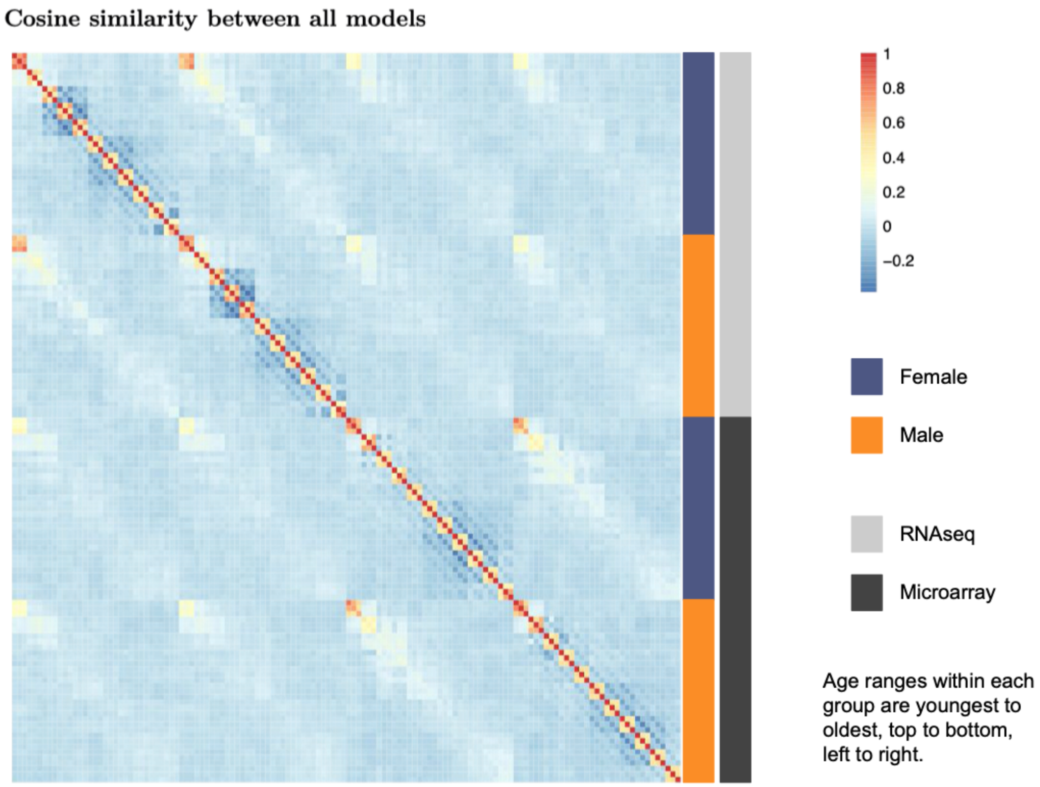


**Figure S22. Cosine similarity of Female and Male model weights.** The heatmap shows the similarity between all RNAseq and microarray models trained in Males. The number after the age range is the number of positive training examples.

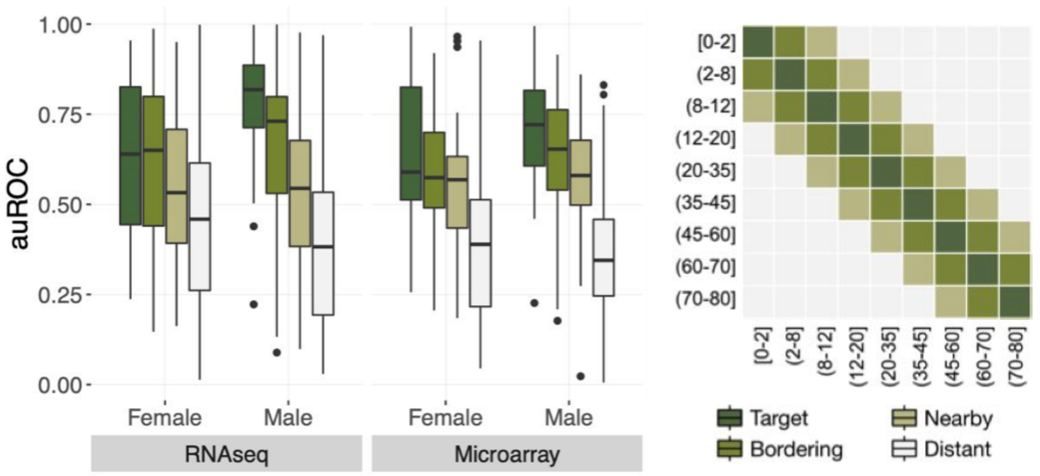


**Figure S23. Performance of blood sample models when evaluated on near and distant age ranges.** Boxplot of auROC performance of all RNAseq and microarray Female and Male models when considering target, bordering, nearby, and distant age ranges as positive examples (key on right).

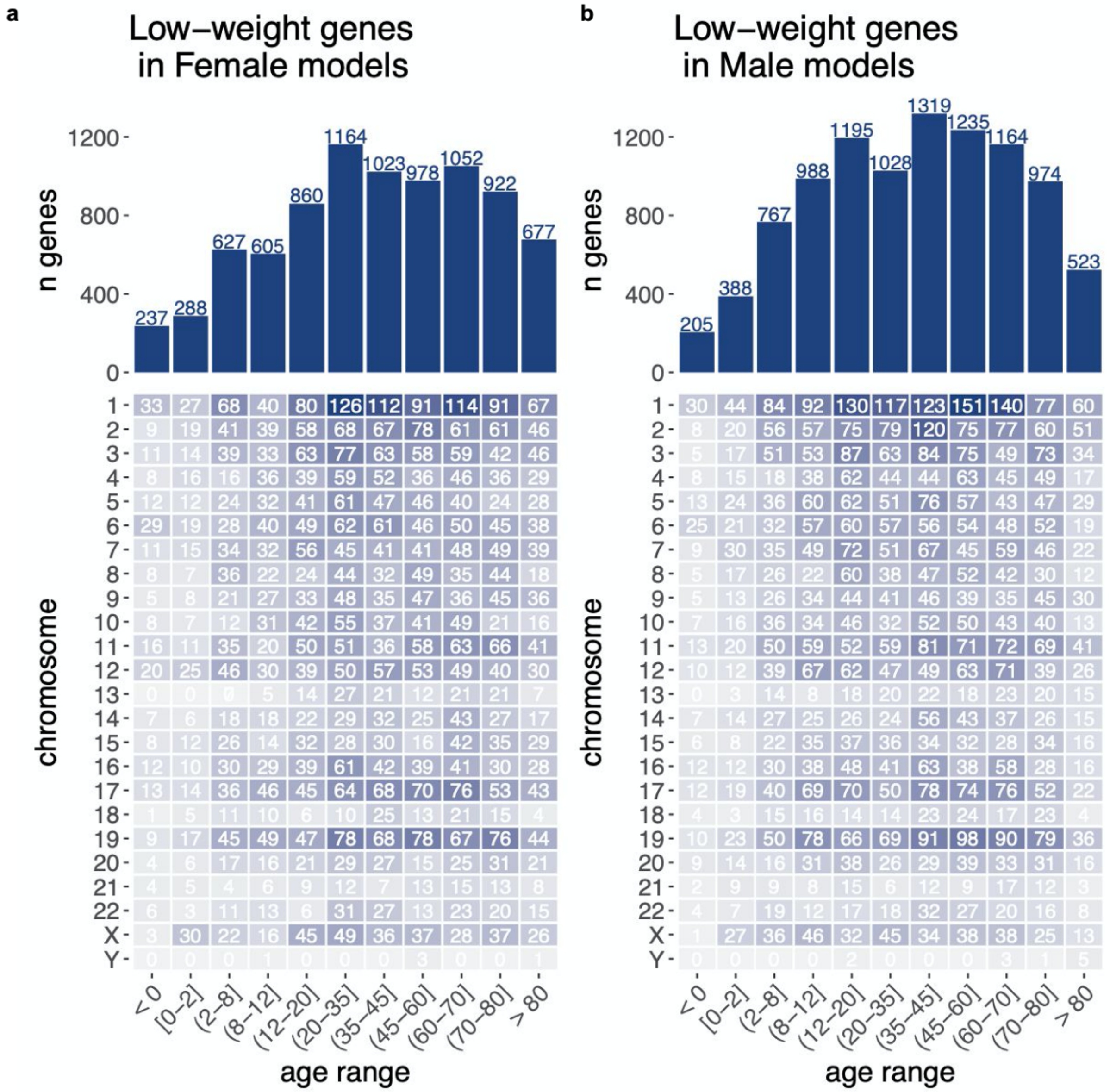


**Figure S24. Number of age-biased genes across age ranges in each sex.** (**a**) The table displays the number of age-biased genes on each chromosome per age range in Females and the barplot shows the total number of age-biased genes per age range. (**b**) The table displays the number of age-biased genes on each chromosome per age range and the barplot shows the total number of age-biased genes per age range in Males.

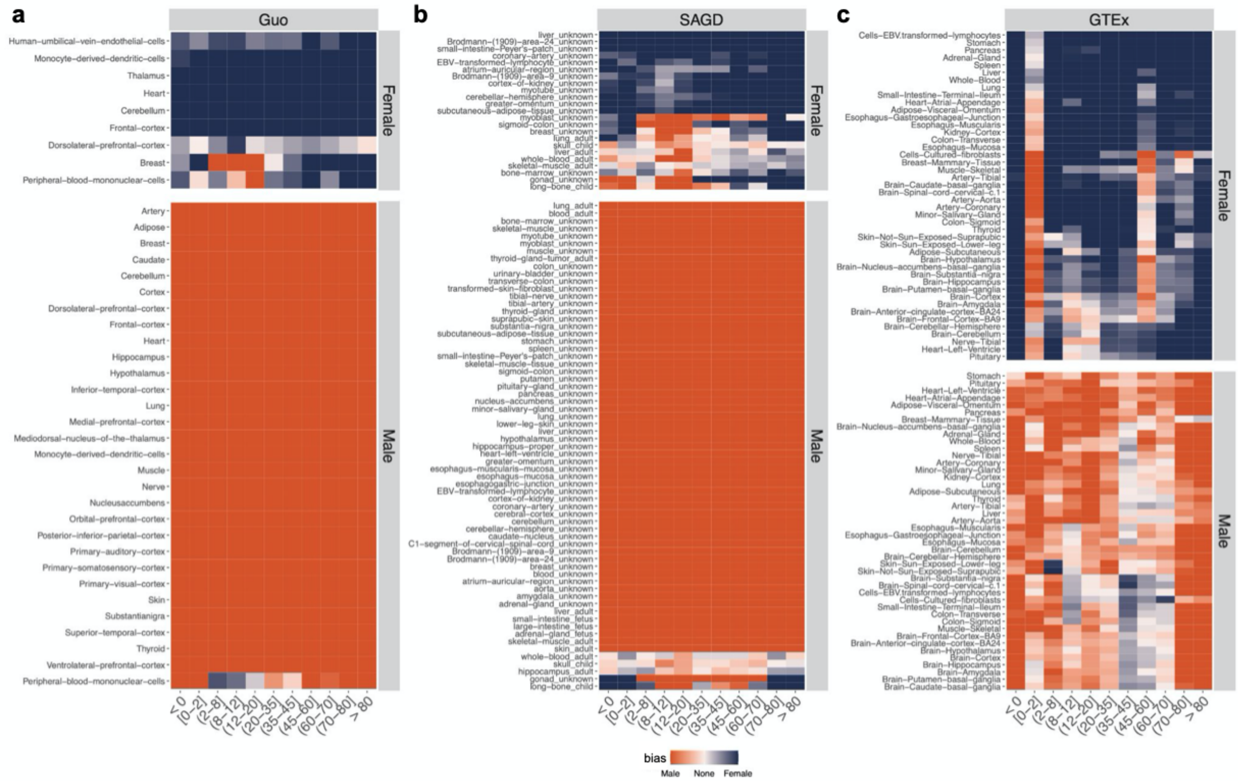


**Figure S25. Enrichment of sex-differentially expressed genes sets from previous studies in our age-stratified sex signatures.** Female- and Male-bias enrichment of gene sets from previous studies. Heatmaps show enrichment scores for (**a**) Guo et al gene sets, (**b**) SAGD gene sets and (**c**) GTEx gene sets.

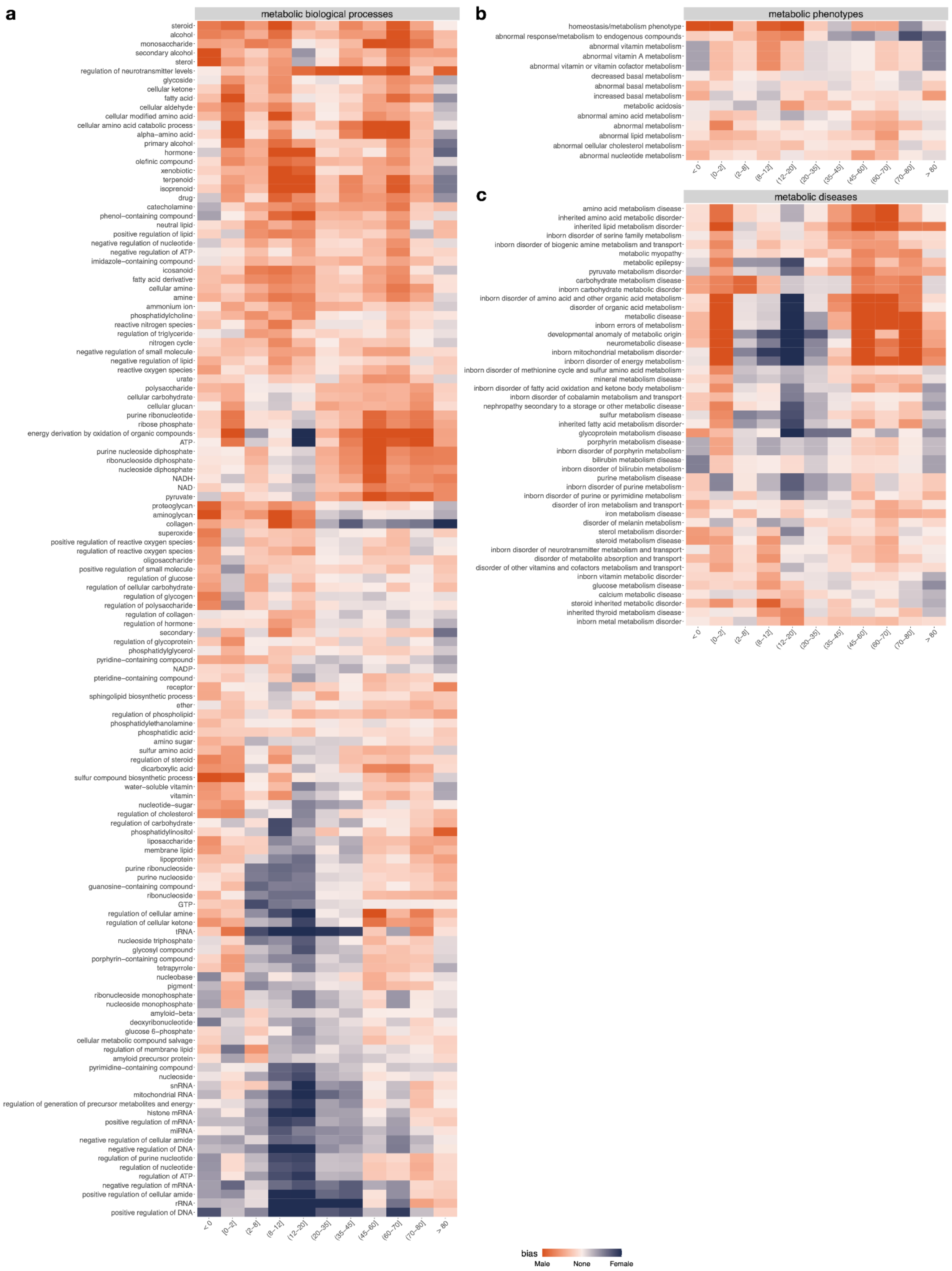

a
metabolic biological processes
steroid
alcohol
monosaccharide
secondary alcohol
sterol
regulation of neurotransmitter levels
glycoside
cellular ketone
fatty acid
cellular aldehyde
cellular modified amino acid
cellular amino acid catabolic process
alpha−amino acid
primary alcohol
hormone
olefinic compound
xenobiotic
terpenoid
isoprenoid
drug
catecholamine
phenol−containing compound
neutral lipid
positive regulation of lipid
negative regulation of nucleotide
negative regulation of ATP
imidazole−containing compound
icosanoid
fatty acid derivative
cellular amine
amine
ammonium ion
phosphatidylcholine
reactive nitrogen species
regulation of triglyceride
nitrogen cycle
negative regulation of small molecule
negative regulation of lipid
reactive oxygen species
urate
polysaccharide
cellular carbohydrate
cellular glucan
purine ribonucleotide
ribose phosphate
energy derivation by oxidation of organic compounds
ATP
purine nucleoside diphosphate
ribonucleoside diphosphate
nucleoside diphosphate
NADH
NAD
pyruvate
proteoglycan
aminoglycan
collagen
superoxide
positive regulation of reactive oxygen species
regulation of reactive oxygen species
oligosaccharide
positive regulation of small molecule
regulation of glucose
regulation of cellular carbohydrate
regulation of glycogen
regulation of polysaccharide
regulation of collagen
regulation of hormone
secondary
regulation of glycoprotein
phosphatidylglycerol
pyridine−containing compound
NADP
pteridine−containing compound
receptor
sphingolipid biosynthetic process
ether
regulation of phospholipid
phosphatidylethanolamine
phosphatidic acid
amino sugar
sulfur amino acid
regulation of steroid
dicarboxylic acid
sulfur compound biosynthetic process
water−soluble vitamin
vitamin
nucleotide−sugar
regulation of cholesterol
regulation of carbohydrate
phosphatidylinositol
liposaccharide
membrane lipid
lipoprotein
purine ribonucleoside
purine nucleoside
guanosine−containing compound
ribonucleoside
GTP
regulation of cellular amine
regulation of cellular ketone
tRNA
nucleoside triphosphate
glycosyl compound
porphyrin−containing compound
tetrapyrrole
nucleobase
pigment
ribonucleoside monophosphate
nucleoside monophosphate
amyloid−beta
deoxyribonucleotide
glucose 6−phosphate
cellular metabolic compound salvage
regulation of membrane lipid
amyloid precursor protein
pyrimidine−containing compound
nucleoside
snRNA
mitochondrial RNA
regulation of generation of precursor metabolites and energy
histone mRNA
positive regulation of mRNA
miRNA
negative regulation of cellular amide
negative regulation of DNA
regulation of purine nucleotide
regulation of nucleotide
regulation of ATP
negative regulation of mRNA
positive regulation of cellular amide
rRNA
positive regulation of DNA
< 0
(0–2]
(2–8]
(8–12]
(12–20]
(20–35]
(35–45]
(45–60]
(60–70]
(70–80]
> 80
b
metabolic phenotypes
homeostasis/metabolism phenotype
abnormal response/metabolism to endogenous compounds
abnormal vitamin metabolism
abnormal vitamin A metabolism
abnormal vitamin or vitamin cofactor metabolism
decreased basal metabolism
abnormal basal metabolism
increased basal metabolism
metabolic acidosis
abnormal amino acid metabolism
abnormal metabolism
abnormal lipid metabolism
abnormal cellular cholesterol metabolism
abnormal nucleotide metabolism
< 0
(0–2]
(2–8]
(8–12]
(12–20]
(20–35]
(35–45]
(45–60]
(60–70]
(70–80]
> 80
c
metabolic diseases
amino acid metabolism disease
inherited amino acid metabolic disorder
inherited lipid metabolism disorder
inborn disorder of serine family metabolism
inborn disorder of biogenic amine metabolism and transport
metabolic myopathy
metabolic epilepsy
pyruvate metabolism disorder
carbohydrate metabolism disease
inborn carbohydrate metabolic disorder
inborn disorder of amino acid and other organic acid metabolism
disorder of organic acid metabolism
metabolic disease
inborn errors of metabolism
developmental anomaly of metabolic origin
neurometabolic disease
inborn mitochondrial metabolism disorder
inborn disorder of energy metabolism
inborn disorder of methionine cycle and sulfur amino acid metabolism
mineral metabolism disease
inborn disorder of fatty acid oxidation and ketone body metabolism
inborn disorder of cobalamin metabolism and transport
nephropathy secondary to a storage or other metabolic disease
sulfur metabolism disease
inherited fatty acid metabolism disorder
glycoprotein metabolism disease
porphyrin metabolism disease
inborn disorder of porphyrin metabolism
bilirubin metabolism disease
inborn disorder of bilirubin metabolism
purine metabolism disease
inborn disorder of purine metabolism
inborn disorder of purine or pyrimidine metabolism
disorder of iron metabolism and transport
iron metabolism disease
disorder of melanin metabolism
sterol metabolism disorder
steroid metabolism disease
inborn disorder of neurotransmitter metabolism and transport
disorder of metabolite absorption and transport
disorder of other vitamins and cofactors metabolism and transport
inborn vitamin metabolic disorder
glucose metabolism disease
calcium metabolic disease
steroid inherited metabolic disorder
inherited thyroid metabolism disease
inborn metal metabolism disorder
< 0
(0–2]
(2–8]
(8–12]
(12–20]
(20–35]
(35–45]
(45–60]
(60–70]
(70–80]
> 80
bias
Male
None
Female

**Figure S26. Enrichment of experimentally-derived genes sets from in our age-stratified sex signatures.** Female- and Male-bias enrichment of experimentally-derived gene sets. Heatmaps show enrichment scores for (**a**) a representative set of metabolism-related GO biological processes, (**b**) metabolic diseases, and (**c**) metabolic phenotypes.
